# Genome-enabled stilbenoid gene cluster discovery in *Pleuropterus multiflorus*

**DOI:** 10.1101/2024.12.13.628289

**Authors:** Jinzhu Jiang, Xianju Liu, Yi Wang, Zhenchang Liang, An Liu

## Abstract

Stilbenoids, a class of compounds with the stilbene skeleton as part of plant secondary metabolism, are renowned for their diverse health benefits. Among them, 2,3,5,4′-Tetrahydroxystilbene-2-*O*-*β*-D-glucoside (THSG), specifically abundant in the famous medicinal plant *Pleuropterus multiflorus*, exhibits significant pharmacological properties. A selfing progeny of *P. multiflorus* commercial variety ‘Jinwufugui No.1’, which contains high THSG content, was used for genome sequencing. Through the combination of weighted correlation network analysis, genome mining, and enzymatic characterization studies, we identified an unpredicted biosynthetic gene cluster responsible for THSG biosynthesis. This cluster includes a stilbene synthase (PmSTS1), a flavin-containing monooxygenase (PmFMO3), and a UDP-glycosyltransferase (UGT72B90). Our findings suggest this stilbenoid gene cluster is formed by transposable element-mediated gene duplications and neofunctionalization, and regulated by MYB and NAC type transcription factors. Notably, PmFMO3, as the first identified stilbenoid-2-hydroxylase, was confirmed that it can modify the other stilbenoids without the A ring modifications. The novel type of gene cluster, in which FMO serves as the key decorating enzyme, will guide the identification of more biosynthetic gene clusters in plants. In addition, the characterization of PmFMO3 extends our understanding of the FMO gene function in plants, and provides an important catalytic bioparts for the heterologous synthesis of polyhydroxylated stilbenoids.

## 1. Introduction

Stilbenoids, a class of compounds with the stilbene skeleton synthesized in plants, play a vital role in their defense mechanism against biotic and abiotic stress [1]. A total of 459 natural stilbenoids have been identified across 45 plant families and 196 plant species until 2022 [2]. They are renowned for their diverse health benefits, such as anti-microbial, anti-cancer, and anti-inflammatory properties as well as protection against heart disease, Alzheimer’s disease, and diabetes [3]. Among stilbenoids, resveratrol is the most popular and extensively studied for its health properties. However, its clinical applications are limited by poor solubility [4] and low bioavailability [5]. Thus, the studies regarding the biological properties of resveratrol derivatives have increased in recent years, and many of them present positive effects. However, limited to the low content, the decorating enzymes that synthesize these resveratrol derivatives are rarely identified in plants, especially for hydroxylases.

The early biosynthetic pathway of stilbenoids is relatively conservative. Stilbene synthase (STS) is the key enzyme, which catalyzes the condensation of three malonyl-CoA and one *p*-coumaroyl-CoA to generate resveratrol6. The catalytic mechanism of STS has been elucidated, and the A ring of resveratrol is formed by the condensation and cyclization of three malonyl-CoA [7]. Both the C-3 and C-5 position hydroxyl groups of the A ring come from the reduction of carbonyl groups [8,9]. Therefore, the other hydroxyl group of A ring should be formed by site-specific hydroxylase.

*Pleuropterus multiflorus* is a famous traditional medicinal plant widely used in East Asia. Its dried roots and caulis have been employed in herbal medicines for over 1000 years [10]. They are rich in stilbenoids, which mainly exist as 2,3,5,4′-tetrahydroxystilbene-2-*O*-*β*-D-glucoside (THSG) [11]. The modern medical studies have confirmed the pharmacological activities of THSG [12–14] and that their content determines the efficacy of the different medicinal parts of *P. multiflorus* [15]. Rare in other stilbenoid-containing plants, the content of THSG in *P. multiflorus* can be up to 9% dry weight in the roots, which makes it an ideal material to identify the decorating enzymes of resveratrol. The identification of THSG biosynthetic pathway will provide the stilbenoid-2-hydroxylase and glycosyltransferase for the heterologous synthesis of stilbenoids (Fig. 1(a)).

**Fig. 1.**
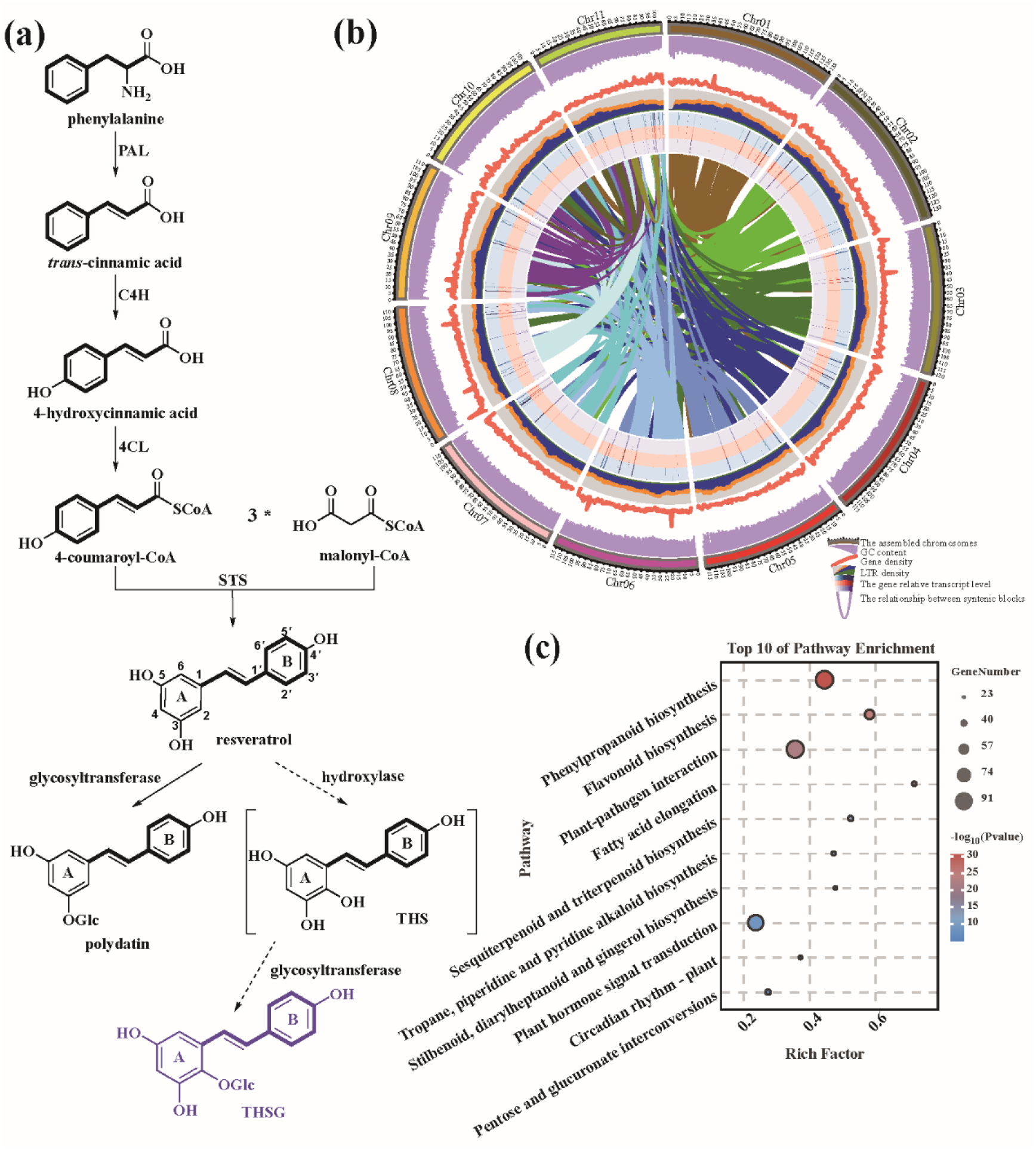
Proposed THSG biosynthetic pathway and the overview of the genomic features of *Pleuropterus multiflorus*. **(a)** Proposed THSG biosynthetic pathway in *P. multiflorus*. Abbreviation: PAL, phenylalanine ammonia lyase; C4H, cinnamate 4-hydroxylase; 4CL, 4-coumarate: coenzyme A ligase; STS, stilbene synthase; THS, 2,3,5,4′-tetrahydroxystilbene; THSG, 2,3,5,4′-tetrahydroxystilbene-2-*O*-*β*-D-glucoside. **(b)** The genomic landscape of the 11 *P. multiflorus* pseudochromosomes. **(c)** Top 10 of pathway enrichment of family expansion genes in *P. multiflorus* by KEGG.

With the development of sequencing in recent years, whole-genome sequencing has become a practical strategy to identify the natural product biosynthetic pathway. A growing number of biosynthetic gene clusters have been identified in plants [16,17]. They are involved in the biosynthesis of terpenoids [18], alkaloids [19], fatty acids [20], benzenoids [21], cyanogenic glycosides [22], flavonoids [23], and phenolamides [24,25]. However, no stilbenoid gene cluster has been identified thus far.

Here, an unpredicted type of gene cluster containing an STS, a flavin-containing monooxygenase (FMO), and a UDP-glycosyltransferase (UGT) was identified to be involved in the THSG biosynthesis by genome sequencing and mining, *in vivo* assays in *Nicotiana benthamiana*, and *in vitro* enzymatic reactions. Resveratrol was biosynthesized by PmSTS1, then its C-2 position was selectively hydroxylated by PmFMO3, finally rapidly glycosylated to product THSG by UGT72B90. This stilbenoid gene cluster was formed by transposable element (TE)-mediated gene duplications and neofunctionalization. As the first identified stilbenoid-2-hydroxylase, the substrate selectivity of PmFMO3 was also identified. The gene cluster expression was positively regulated by MYB and NAC type transcription factors. The gene cluster of THSG biosynthesis reveals a novel type of gene cluster in plants, in which FMO serves as the key decorating enzyme, and also can guide to elucidate the biosynthetic pathways of other bioactive compounds in plants. The characterization of PmFMO3 provides an important catalytic bioparts for the heterologous synthesis of polyhydroxylated stilbenoids. In addition, the characterization of THSG biosynthetic pathway provides important reference information for germplasm evaluation and molecular breeding of *P. multiflorus* based on bioactive compounds.

## 2. Materials and methods

### 2.1. Plant materials and growth conditions

The plant for genome sequencing is a selfing progeny of *P. multiflorus* commercial variety ‘Jinwufugui No.1’, which was bred for high THSG contents and widely cultivated in Yunnan Province. It was planted in a greenhouse at 25°C under a light period of 16-h light/8-h dark. The fresh leaves were collected to exact high-quality genomic DNA for genome sequencing. For transcriptome sequencing and THSG content quantification, three-year-old cultivated *P. multiflorus* ‘Jinwufugui No.1’ were collected and separated into 15 different tissues, including buds (B), bark of caulis (BC), cutting stem (CS), expanded root tubers (ERT), flowers (F), mature leaves (ML), the petioles of mature leaves (PML), rachis (R), seeds (S), thick brown lignified caulis (d ≥ 4 mm) (TcBLC), thin brown lignified caulis (d < 4 mm) (TnBLC), unexpanded root tubers (URT), young green caulis (YGC), xylem of caulis (XC), and young leaves (YL). To perform transient expression assays in tobacco leaves, the *N. benthamiana* were grown in the greenhouse at 25°C under a light period of 16-h light/8-h dark.

### 2.2. Genome sequencing, assembly, and annotation

The genome sequencing, assembly, and annotation of *P. multiflorus* were processed with the previously described method [26], and the detailed methods were listed in the Supplementary methods.

### 2.3. Genome evolution analysis

To clarify the phylogenetic relationships of *P. multiflorus*, 11 additional species were selected for the analysis, including *Fagopyrum esculentum*, *F. tataricum*, *Rumex hastatulus*, *Vitis vinifera*, *Senna tora*, *Arachis duranensis*, *A. hypogaea*, *Morus notabilis*, *Populus trichocarpa*, *Theobroma cacao*, and *Oryza sativa* (Table S1). The orthologous genes among all 12 species were clustered by OrthoMCL [27]. Single-copy genes were extracted from the clustering results and multiple sequence alignments were performed by MUSCLE [28]. High-quality aligned protein sequences remained after removing low-quality alignment or divergent regions by Gblocks [29]. The phylogenetic tree was constructed by RAxML with the PROTGAMMAJTTF model and bootstrap of 1000 [30]. MCMCtree from the PAML package [31] was used to estimate the species divergence time according to TimeTree (http://www.timetree.org). Three divergence times were used in this analysis, including those of *T. cacao* and *P. trichocarpa*, *V. vinifera* and *P. trichocarpa*, and *V. vinifera* and *O. sativa*. The OrthMCL results and time divergence tree were used as the input for the CAFÉ program, which was used to identify expansions and contractions of gene families across 12 plant genomes. Expanded orthogroups were defined according to *P*-value < 0.05 and a gene number greater than the average value of multiple species.

### 2.4. Genome synteny and whole-genome duplication (WGD) event analysis

All-vs-all BLASTP [32] (*E*-value 1e−05) and MCScanX [33] were used to predict the collinear relationships and positional features between *P. multiflorus* and other species. Blocks of >10 genes and gaps of <5 genes were obtained. The synteny map and dot plot were processed by MCScan and drawn by the python scripts in MCScan packages [34].

The segment duplication events were predicted using self-vs-self BLASTP [32] (*E*-value 1e−05) and MCScanX [33] among the *P. multiflorus* genome, requiring at least five genes per collinear block. Subsequently, the pairwise sequences from the synteny blocks and segment duplication pairs were processed by ParaAT [35]. The values of synonymous substitutions per site (Ks) were calculated by Kaks_Calculator [36], and the distribution of Ks within paralogs was used to examine the most recent WGD event in *P. multiflorus* genome.

### 2.5. mRNA-sequencing and transcriptome analysis

Total RNA was extracted from the different tissue samples using an RNAprep Pure Plant Kit (TIANGEN, China). Then the libraries for mRNA-sequencing were constructed by Biomarker Technologies and sequenced using the Illumina NovaSeq 6000 platform. The raw mRNA-sequencing data were trimmed with TRIMMOMATIC [37]. The clean reads were mapped to genome assembly by TopHat [38] with default parameters. The expression levels of genes represented by FPKM (fragments per kilobase million) for each sample were calculated by Cufflink [38] with default parameters.

### 2.6. Quantification of THSG by UPLC-DAD

The THSG contents of *P. multiflorus* different tissues were quantified by ultra-high-performance liquid chromatography coupled with diode array detector (UPLC-DAD). 25 mL extraction buffer containing 50% ethanol was added to 0.75 g frozen plant powder. After ultrasonic treatment (650W, 40Hz) for 30 min and centrifugation at 13,000 × g for 5min, the supernatant was used for UPLC-DAD analysis by Thermo Scientific Vanquish Flex UPLC-DAD system. 2 μL volume of each extract was analyzed using a Waters ACQUITY UPLC BEH C18 column (2.1 mm× 150 mm, 1.7 μm), and the column was maintained at 40℃. The mobile phases were 0.1% formic acid aqueous solution (A) and acetonitrile (B), respectively. The elution gradient was performed as follows: 0.3 mL/min flow rate; 0 min 10% B; 8 min 18% B; 14 min 22% B; 19 min 33% B; 22 min 90% B; 26 min 90% B; 26.1 min 10% B; 30 min 10% B. *trans*-THSG was detected at 320 nm. The THSG reference standards obtained from China National Institutes for Food and Drug Control was used to identify elution times and accurate quantification.

### 2.7. Gene coexpression network and cluster analysis

To identify the relationships between the expressed genes, weighted correlation network analysis (WGCNA) [39] was performed. A signed coexpression network was constructed using a soft-thresholding power of eight and default parameters. Finally, we obtained 12 clusters for these genes.

### 2.8. Genome-wide identification of CYP450, FMO, and UGT gene families in *P. multiflorus*

The conserved CYP450, FMO, and UGT domain sequence (Pfam ID: PF00067, PF00743, and PF00201) were used to identify all possible CYP450, FMO, and UGT protein candidates in the *P. multiflorus* genome by simple Hidden Markov Model (HMM) search of TBtools [40], respectively. In addition, the *Arabidopsis thaliana* CYP450, FMO, and UGT gene family proteins were used as the queries to identify the corresponding candidates in the *P. multiflorus* genome by BLASTP [32] (*E* < 10^−5^). The candidates obtained from these two methods were combined as the corresponding family candidates, and NCBI-Conserved Domain Database (CDD) [41] search was processed to confirm they contained the corresponding family characteristic domain. And the sequences without the signature motifs (CYP450: the PERF consensus (PXRX) and the K-helix (XEXXR); FMO: FAD binding motif (GXGXXG), ‘TGY’ motif, and FMO-identifying motif (FXGXXXHXXXY/F) and UGT: PSPG motif) were removed. The *A. thaliana* proteins were used as the comparative method to align with the predicted gene family members by ClustalW [42] program. The phylogenetic tree was constructed using maximum likelihood (ML) or neighbor joining (NJ) method by MEGA11 [43]. The CYP450 and UGT family members were officially named by the International P450 Nomenclature Committee and UGT Nomenclature Committee, respectively. The FMO family members were named based on their chromosome location.

### 2.9. The enzyme activity assay in *N. benthamiana* leaves

To evaluate the activity of the candidate enzymes, they were transiently over-expressed in *N. benthamiana* leaves, respectively or jointly. Firstly, the full-length CDSs of *CYP450s*, *PmFMOs*, *UGTs*, and *PmSTS1* were cloned into the transient expression vector pEAQ-HT-DEST using GATEWAY technology to generate the corresponding destination vectors, respectively. Subsequently, they were independently transformed into *Agrobacterium tumefaciens* GV3101. The overnight cultured engineered *A. tumefaciens* were centrifuged, and the pellets were resuspended in infiltration buffer containing 10 mM 4-morpholineethanesulfonic acid (MES, pH 5.6), 10 mM MgCl_2_, and 150 mM acetosyringone to an OD_600_ of 0.5. Tobacco leaves were infiltrated with the resuspension by using the 1 mL needleless syringe. Three days later, tobacco leaves were infiltrated with the substrate solution containing 50 μM resveratrol or other stilbenoids. Finally, the tobacco leaves infiltrated with substrates were harvested after 24 h cultivation and lyophilized for metabolite analysis. Each group had three biological replicates. The primers used for gene cloning and constructing overexpression plasmids are listed in Table S2.

### 2.10. *In vitro* enzyme activity assay

To characterize the activity of FmFMO3 and UGT72B90 *in vitro*, we cloned their coding sequence from the cDNA of *P. multiflorus* and inserted them into the prokaryotic expression vector pET32a. The NAD(P)H: flavin oxidoreductase (Fre) was cloned into pET28a as previous report [44]. The vectors were transformed into *Escherichia coli* BL21 competent cells. The expression of the His-fusion proteins was induced with 0.1 mM IPTG at 16 °C for 24 h. The recombinant proteins were purified by Qiagen Ni-NTA resin. To detect the activity of the proteins, a 500 μL enzyme reaction system comprises 50 mM Tris-HCl buffer (pH 7.5), 100 μM resveratrol or other stilbenoids, 20 μM FAD, 2 mM NADPH, 20 μg Fre, and 100 μg FmFMO3 or GUS protein were utilized. 500 μM UDP-Glc and 100 μg UGT72B90 were also added to the reaction system when glycosyltransferase activity was detected. The enzyme assays were incubated at 30°C for 24 h and quenched with equivalent methanol to precipitate enzymes. All assays were repeated three times. The primers used for their cloning and constructing prokaryotic expression plasmids are listed in Table S2.

### 2.11. Characterization and quantification of the stilbenoids by UPLC-Q-TOF-MS and UPLC-QQQ-MS

We detected the stilbenoids by an Agilent UPLC system (1290 infinity 2 series) with an Agilent 6540 Q-TOF mass spectrometer equipped with an Agilent Jet Stream Source in negative ion mode and auto MS/MS mode. The UPLC parameters were the same as those used in THSG Quantification. The source parameters were as follows: drying gas (N_2_) temperature and flow rate, 300℃ and 9.0 L min^−1^; nebulizer, 35 psig; sheath gas temperature and flow rate, 400℃ and 11.0 L min^−1^; Vcap, 3500 V; skimmer, 65 V; OCT 1 RF Vpp, 750 V; fragmentor voltage, 120 V, and the collision energy for MS/MS was 35 V.

After the stilbenoids characterization, one precursor ion and two corresponding characteristic ions of each stilbenoid were chosen to quantify them by an Agilent UPLC system (1290 infinity 2 series) with an Agilent G6460A triple quadrupole mass spectrometer equipped with an Agilent Jet Stream Source in negative ion mode and multiple reaction monitoring (MRM) mode. The source parameters were as follows: drying gas (N_2_) temperature and flow rate, 300℃ and 8.0 L min^−1^; nebulizer, 30 psig; sheath gas temperature and flow rate, 300 ℃ and 11.0 L min^−1^; capillary, 3500 V. The parameters of fragmentor and collision energy for each stilbenoid were listed in Table S3.

### 2.12. Luciferase assay

To evaluate the regulatory role of the candidate transcription factors to the gene cluster, the full-length CDSs of these transcription factors were cloned into the transient expression vector pEAQ-HT-DEST using GATEWAY technology to generate the corresponding destination vectors, respectively. The 2k upstream regions of *PmSTS1*, *UGT72B90*, and *PmFMO3* were cloned into the pGreen0800-LUC vector by homologous recombination. Then they were independently transformed into *Agrobacterium tumefaciens* GV3101 carrying the helper plasmid pSoup-P19. The overnight cultured positive clones were centrifuged, and the pellets were resuspended in infiltration buffer to an OD_600_ of 0.6. Then the resuspensions harbor pEAQ and pGreen0800-LUC vector were combined in a 2:1 ratio, and infiltrated into the tobacco leaves. six days later, the tobacco leaves were infiltrated with luciferase assay reagent, and visualized with a chemiluminescence instrument. The primers used for gene and promoter cloning and constructing plasmids are listed in Table S2.

## 3. Results

### 3.1. Genome sequencing, assembly, annotation, and phylogenomic analyses

*P. multiflorus* commercial variety ‘Jinwufugui No.1’ was bred for high THSG contents (greater than 2% in flesh root tuber), and had been widely cultivated in Yunnan Province. A selfing progeny was chosen for genome sequencing. A 1.3 Gb high-quality chromosomal-level reference genome sequence of *P. multiflorus* was obtained by the combination of PacBio HiFi and Illumina data and assembled using information from Hi-C technologies (Fig. 1(b)). The LAI (LTR Assembly Index) score was 19.35, reaching the criterion of reference quality. The high assembled quality provided the basis for subsequent analyses. The information about the genome statistics and annotation was listed in Tables S4-10 and shown in Figs. S1, S2.

The annotated genes were clustered into gene families among *P. multiflorus* and the other 11 species, including three Polygonaceae species (*Fagopyrum esculentum*, *F. tataricum*, and *Rumex hastatulus*), five stilbenoid-containing species (*Vitis vinifera*, *Senna tora*, *Arachis duranensis*, *A. hypogaea*, and *Morus notabilis*), and three other angiosperms (*Populus trichocarpa*, *Theobroma cacao*, and *Oryza sativa*). We selected 201 single/low-copy gene families among these 12 species to construct the phylogenetic tree, which showed that *P. multiflorus* and *R. hastatulus* belong to a clade, while *F. esculentum* and *F. tataricum* belong to the other clade. Based on molecular clock analysis, we estimated that these two clades diverged ∼82 million years ago (Mya) (55–111 Mya), and *P. multiflorus* diverged from *R. hastatulus* ∼69 Mya (44–99 Mya) (Fig. S3). We conducted expansion and contraction analysis based on the constructed phylogenetic tree and discovered 1212 expanded and 53 contracted families in *P. multiflorus* (Fig. S3). The KEGG functional enrichment analysis of the expanded gene families demonstrated that they were mainly assigned to “Phenylpropanoid biosynthesis”, “Flavonoid biosynthesis”, and “Plant-pathogen interaction” (Fig. 1(c)).

By constructing the distribution of synonymous substitution rates per gene (K_S_) within *P. multiflorus* and *F. tataricum*, three polyploidization events were detected in the genome of *P. multiflorus* (Fig. S4). The distribution of the reciprocal best hit (RBH) gene pair K_S_ values exhibited a peak at ∼2.05 in these two species, representing the ancient core eudicot *γ* triplication event [45]. Whereas the second prominent Ks peak at 0.79 existed in these two species, reflecting one WGD event common to all Polygonaceae species. But the Ks peak at 0.79 was asymmetrical, there seemingly exist another peak. Then we selected the big segments which contained more than 50 genes to re-construct the distribution of K_S_, and found another peak at 0.70 after its split from *F. tataricum*. Synteny analyses between the genomes of *P. multiflorus*, *F. tataricum*, *Pyrus bretschneideri*, and *V. vinifera* also showed clear evidence of another recent WGD event for *P. multiflorus*. For each genomic region in *V. vinifera*, we typically found four matching regions in *P. multiflorus* with a similar level of divergence and identified 4:2 and 2:2 syntenic depth ratios in the *P. multiflorus*-*Pyrus bretschneideri* and *P. multiflorus*-*F. tataricum* genome comparison, respectively (Figs. S5-7).

The proliferation of repetitive elements, especially the Gypsy or Copia family, was regarded as the main cause for the genome expansion of species in the Polygonaceae family. 883Mb (67.80%) of sequences in the assembled *P. multiflorus* genomes were identified as transposable element (TE), and the predominant type of TE was LTR elements (Tables S9). Among the LTRs, most were Gypsy elements, which accounted for 30.04% of the total repetitive sequences in *P. multiflorus*. We then estimated the times of the LTR burst in the *P. multiflorus* genome, and the results suggested that the timing of the main LTR burst was ∼163.1 Kya, which mainly contributed by Gypsy elements (Fig. S8).

### 3.2. Transcriptome and gene co-expression analysis

To identify the key synthetases involved in the biosynthesis of THSG, the whole plant of *P. multiflorus* was divided into 15 tissues (Fig. 2(a)). All the tissues were used for the transcriptome and THSG content analysis (Fig. 2(b)). Using weighted gene co-expression network analysis (WGCNA) and the dynamic hierarchical tree cut approach, 12 unique modules with comparable gene expression patterns were evaluated (Fig. S9). Among the modules, the eigengene of module blue was highly correlated with the THSG content by Pearson correlation analysis (Fig. 2(c)). Then 28 hub genes which had high gene significance for THSG content (GS > 0.85) and module membership (MM > 0.8) were identified in the module blue (Fig. 2(d); Tables S11). Among them, the hub gene *PM02g9825* was annotated as an *STS* gene. Its protein similarities with reported PmPKS [46] and PmSTS2 [47] were 98.68% and 98.94%, respectively (Fig. S10). We speculated *PmPKS* and *PmSTS2* should be the variants of this chromosomal locus, therefor we named *PM02g9825* as *PmSTS1*. When *PmSTS1* was transiently over-expressed in *N. benthamiana*, *trans*-piceid can be detected as previous report (Fig. S11). Besides *PmSTS1*, three hub genes were annotated as transcription factors (PM02g6581, WRKY transcription factor, PM11g55274, MYB transcription factor, and PM02g10052, NAC transcription factor) (Fig. 2(d)). They may involve in the regulation of THSG biosynthesis in *P. multiflorus*. Unfortunately, no reported type of hydroxylase and glycosyltransferase was found in these hub genes.

**Fig. 2.**
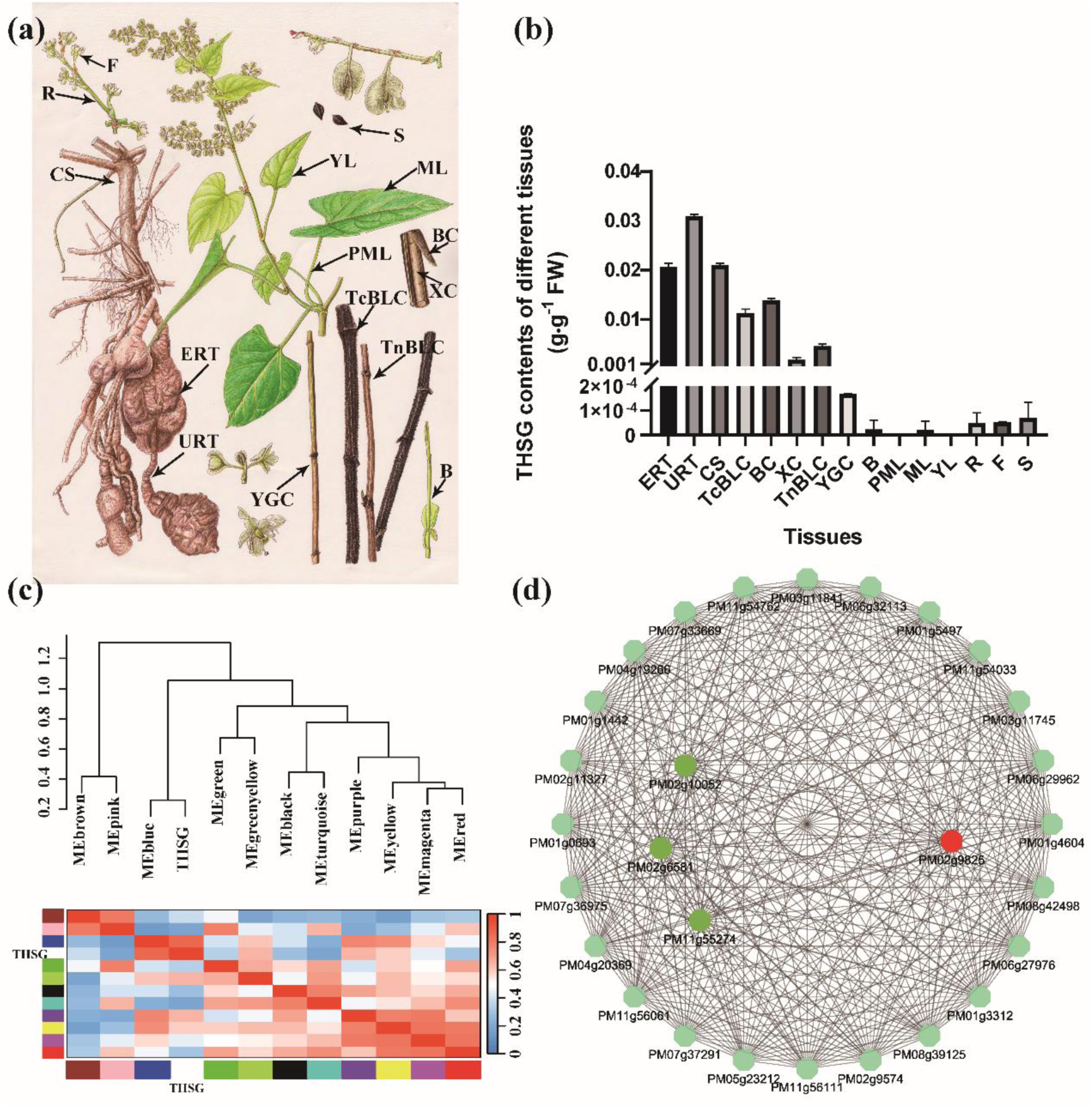
Weighted gene co-expression network analysis (WGCNA) and the evolution of CYP450 and flavin-containing monooxygenase (FMO) gene family in *Pleuropterus multiflorus*. **(a)** Morphology of the 15 tissues of *P. multiflorus* used for RNA-seq and THSG contents. **(b)** The THSG contents of the *P. multiflorus* 15 tissues. B: buds, BC: bark of caulis, CS: cutting stem, ERT: expanded root tubers, F: flowers, ML: mature leaves, PML: the petioles of mature leaves, R: rachis, S: seeds, TcBLC: thick brown lignified caulis (d ≥ 4 mm), TnBLC: thin brown lignified caulis (d < 4 mm), URT: unexpanded root tubers, YGC: young green caulis, XC: xylem of caulis, and YL: young leaves. **(c)** The relationship between THSG content and the modules evaluated by the WGCNA. **(d)** Co-expression network of 28 hub genes identified in the module blue. The red hexagon represents the key stilbene synthesis gene; The green hexagons represent the transcription factors; The pale green hexagons represent the other hub genes.

### 3.3. The identification of stilbenoid-2-hydroxylase in *P. multiflorus*

On account of the reported prokaryotic stilbenoid-3′-hydroxylases belong to heme-containing cytochrome P450 (CYP450) [48–50] and FMO [44,51] families, we identified the CYP450 and FMO gene families in *P. multiflorus* genome based on Hidden Markov model (HMM) and BLASTP search method. Then 115 CYP450 and 28 FMO genes which contain their family signature motifs were found and named, respectively (Fig. 3(a) and (b); Table S12, S13). In the module blue, only two CYP450 genes (CYP81DB2 and CYP71AT253) and one FMO gene (PmFMO3) had relatively high correlation with THSG contents (> 0.70), and the expressions of *CYP71AT253* were low in the THSG-enriching tissues. Interesting, the expressions of two suspected CYP450 genes (PM04g17426 and PM08g40140) in the module blue also had relatively high correlation with THSG content (0.79 and 0.74, respectively). They were annotated as CYP450 by the public databases, but lacked partial family signature motifs (PM04g17426, which lacking PERF and K-helix motif, and PM08g40140, which the C-terminus of CYP450 conserved domain is incomplete). Combining the correlation with THSG content, annotation information, and gene expression level, CYP81DB2, PmFMO3, PM04g17426, and PM08g40140 were chosen as the candidates of stilbenoid-2-hydroxylase. Then their functions were identified by transient expression in *N. benthamiana*. When fed with *trans*-resveratrol, the aglycone of THSG (*m/z* 243.0667 [M-H]^−^) cannot be detected in any transient transgenic leaf. But THSG (*m/z* 405.1198 [M-H]^−^) was remarkably detected in the *PmFMO3*-overexpression leaves, while the *CYP81DB2*, *PM04g17426*, or *PM08g40140*-overexpression leaves showed no difference with the control leaves (Fig. 3(c) and (d)). When *PmSTS1* and *PmFMO3* were simultaneously over-expressed in *N. benthamiana* leaves, THSG still can be obviously detected when no exogenous substrate was added (Fig. S12). To further confirm the function of PmFMO3, the recombinant PmFMO3 protein expressed in the *Escherichia coli* BL21 (DE3) cell and purified by Ni-NTA resin was added to the *in vitro* enzymatic reaction. But neither the aglycone of THSG nor THSG could be detected when *trans*-resveratrol was used as the substrate (Fig. S13). We speculated that the aglycone of THSG may be unstable, and should be stabilized by glycosylation. This speculation was confirmed by the result of computational chemistry, the degradation of the aglycone of THSG was favorable from thermodynamic view (Fig. S14).

**Fig. 3.**
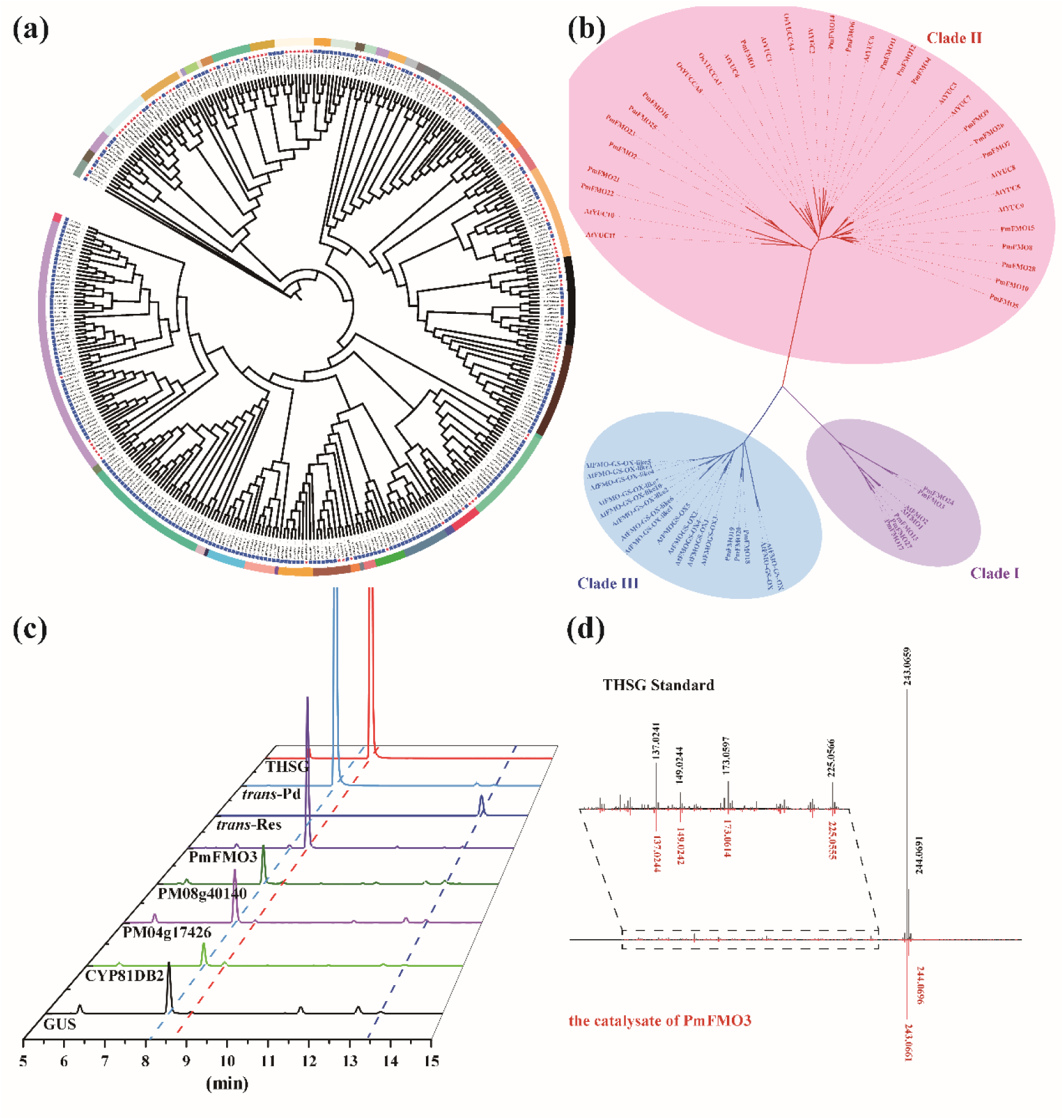
The identification of stilbenoid-2-hydroxylase in *Pleuropterus multiflorus*. **(a)** Phylogenetic tree of CYP450 gene family in *P. multiflorus*. The genes from *P. multiflorus* were marked with red star, and the genes from *A. thaliana* were marked with blue quadrate. Different color in the outlayer represented different subfamilies of CYP450. **(b)** Phylogenetic tree of FMO gene family in *P. multiflorus*. Different color represented different clade of FMO. **(c)** The identification of *P. multiflorus* stilbenoid-2-hydroxylase candidates in *N. benthamiana*. **(d)** Secondary fragment ions alignment between THSG standard and the catalysate of PmFMO3.

### 3.4. The identification of stilbenoid-2-glycosyltransferase in *P. multiflorus*

To identify the glycosyltransferase involving in the biosynthesis of THSG, we also identified the UDP-glycosyltransferase (UGT) gene family in *P. multiflorus* genome based on HMM and BLASTP search method, and found 117 UGT genes containing the plant secondary product glycosyltransferase (PSPG) motif in the whole *P. multiflorus* genome (Fig. 4(a); Table S14). Interesting, we found *UGT72B90* (chr02: 101,190,670-101,192,100) located between *PmSTS1* (chr02: 100,961,857-100,963,850) and *PmFMO3* (chr02: 101,753,609-101,773,663) in the chromosome 02 (Fig. 4(b)). Besides of UGT72B90, two other UGT genes (UGT89X17, and UGT71U19) had relatively high correlation with THSG content (0.75 and 0.64, respectively) and their gene functions were annotated as flavonoid glycosyltransferase. These three UGTs were chosen as the candidate genes of stilbenoid-2-glycosyltransferase and functionally verified by *N. benthamiana* transient expression analysis. When fed with *trans*-resveratrol, the THSG content in *PmFMO3/UGT72B90*-overexpression leaves was significantly more than the control leaves, while *PmFMO3/UGT89X17* and *PmFMO3/UGT71U19*-overexpression leaves showed no difference with the control leaves (Fig. 4(c)). We speculated UGT72B90 was the stilbenoid-2-glycosyltransferase. To further identity the function of UGT72B90, recombinant UGT72B90 protein was expressed in the *E. coli* BL21 (DE3) cell and purified by Ni-NTA resin. When the recombinant PmFMO3 and UGT72B90 proteins, and *trans*-resveratrol were simultaneously added to the *in vitro* enzymatic reaction, THSG could be detected (Fig. 4(d)).

**Fig. 4.**
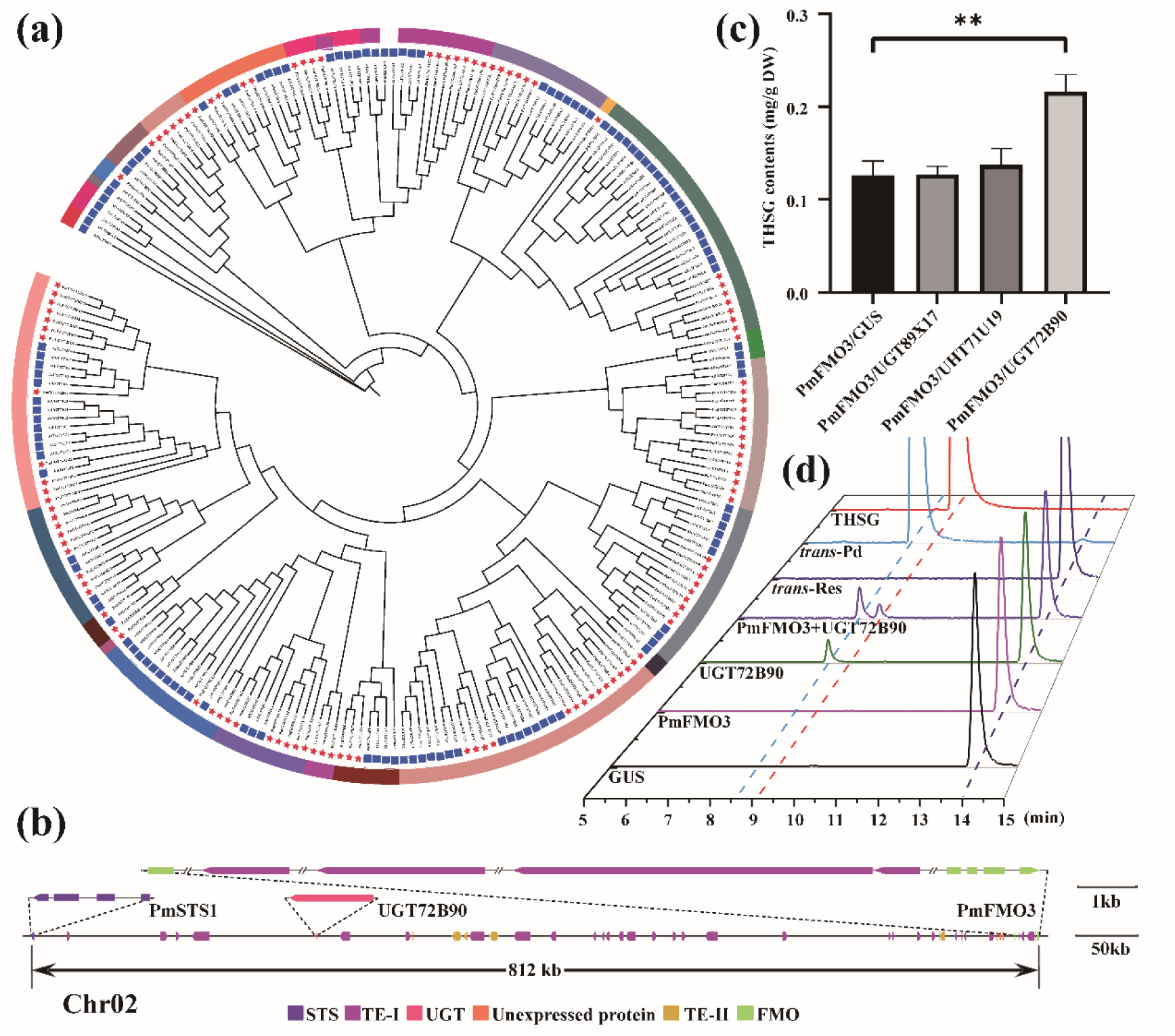
The identification of stilbenoid-2-glycosyltransferase in *Pleuropterus multiflorus*. **(a)** Phylogenetic tree of UDP-glycosyltransferases (UGT) gene family in *P. multiflorus*. The genes from *P. multiflorus* were marked with red star, and the genes from *A. thaliana* were marked with blue quadrate. Different color in the outlayer represented different subfamilies of UGT. **(b)** The display of the sequence information of the section of chromosome 02 that contains *PmSTS1*, *UGT72B90*, and *PmFMO3*. **(c)** The identification of *P. multiflorus* stilbenoid-2-glycosyltransferase candidates in *N. benthamiana*. **(d)** The function characterization of PmFMO3 and UGT72B90 separation or combination *in vitro*.

### 3.5. The discovery of the stilbenoid gene cluster in *P. multiflorus*

Because *PmSTS1*, *UGT72B90*, and *FmFMO3* locate closely in the chromosome 2 (Fig. 4(b)), we speculated they may work as a gene cluster to involve in the THSG biosynthesis in *P. multiflorus*. By analyzing the gene expression information of *PmSTS1*, *PmFMO3*, and *UGT72B90*, we found they shared a very similar expression patterns in *P. multiflorus,* their expressions were significant correlation (*P* < e^−9^) (Fig. 5(a)). We therefore conclude this is a novel type of gene cluster which involves in the complete biosynthesis of THSG, the species-specific stilbenoid in *P. multiflorus*, from the precursor *p*-coumaroyl-CoA and malonyl-CoA. In order to explore the formation mechanism of this gene cluster in *P. multiflorus*, the sequence information of the cluster was re-checked. Many TEs were found to disperse in the cluster, especially four Copia-type retrotransposons were inserted in the first intron of *PmFMO3* (Fig. 4(b)). Therefore, we calculated the TE density, gene density and the ratio of TE/gene density across the section of chromosome 02 that contains this gene cluster (Chr02: 100,961,857-101,773,663), and a higher TE/gene density ratio was observed within the location of the gene cluster in comparison with the adjacent chromosomal regions (Fig. 5(b)). In addition, gene duplication events in *P. multiflorus* were detected, and 45,344 duplicated genes were identified and grouped into five different categories (Table S15). Both *PmFMO3* and *UGT72B90* are transposed duplicates, their ancestral chromosomal locations were *PmFMO13* (PM07g33546) and UGT72B91 (PM05g24154), respectively. Collectively, these results suggested that highly enriched TEs may have brought the gene *PmFMO3* and *UGT72B90* to form a unique gene cluster to produce THSG in *P. multiflorus*.

**Fig. 5.**
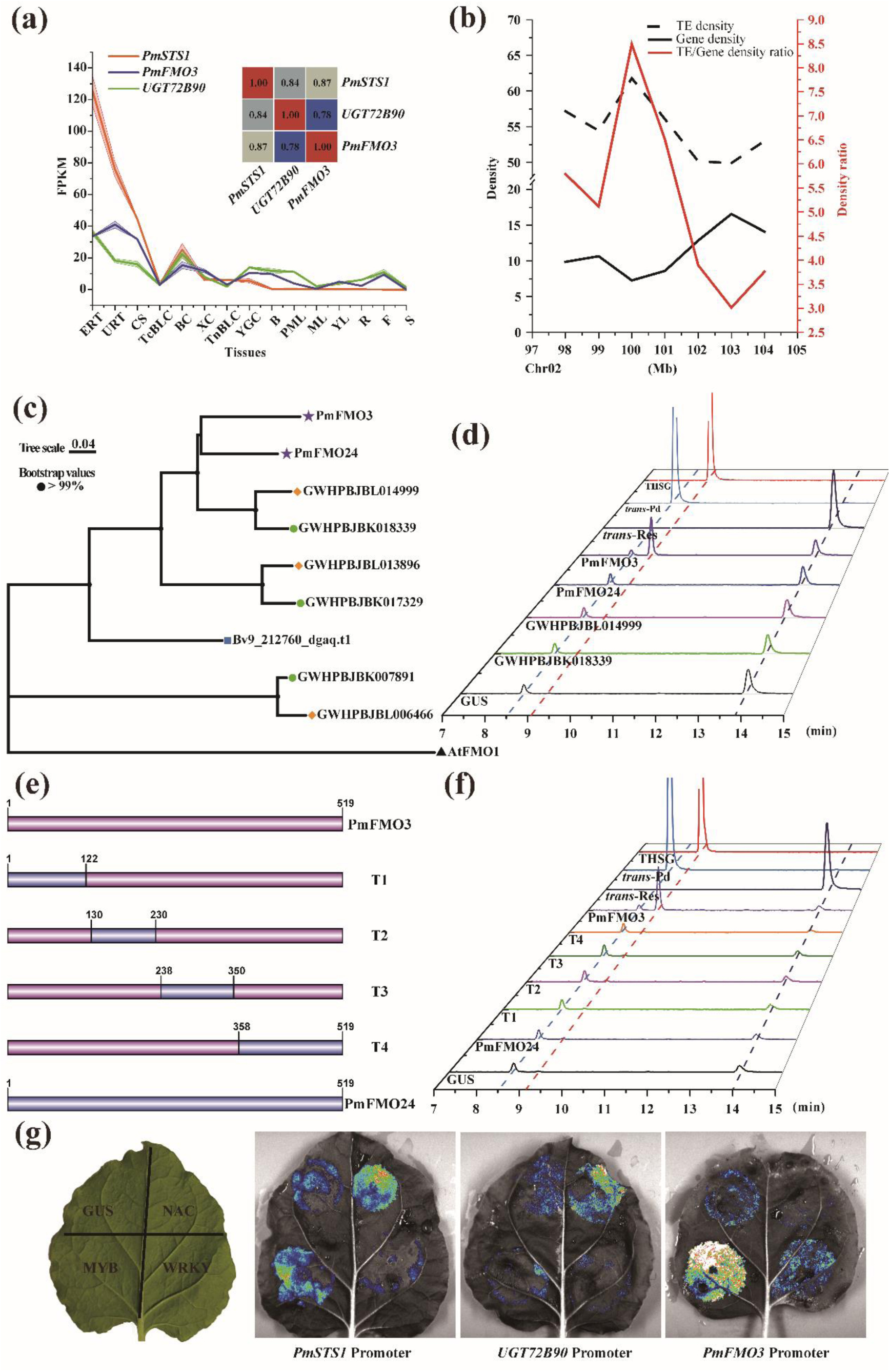
The discovery and formation mechanism of the stilbenoid gene cluster in *Pleuropterus multiflorus*. **(a)** The expression pattern of *PmSTS1*, *PmFMO3*, and *UGT72B90* in 15 tissues of *P. multiflorus*, and the co-expression coefficients between each other. **(b)** Distribution of gene density (black line), TE density (black dashed line), and the TE/gene density ratio (red line) across a section of chromosome02 that contains the stilbenoid gene cluster. **(c)** The phylogenetic tree of some homologous genes of PmFMO3. It was built by MEGA11 with NJ method, and AtFMO1 was used as outgroup. Different colors and shapes represented different plant species. Purple star: *P. multiflorus*, orange diamond: *Fagopyrum tataricum*, green circle: *F. esculentum*, Blue square: *Beta vulgaris*, Green triangle: *Arabidopsis thaliana*. **(d)** The function characterization of PmFMO24, GWHPBJBK018339, and GWHPBJBL014999 in *N. benthamiana* leaves by UPLC-QQQ-MS. *PmFMO24*, *GWHPBJBK018339*, and *GWHPBJBL014999* were cloned from the flower of *P. multiflorus*, the seeds of *Fagopyrum esculentum* and *F. tataricum*, respectively. Then, they were overexpressed in *N. benthamiana* leaves, and fed with resveratrol. PmFMO3 was used as the positive control. **(e)** Schematic picture of the mutations (T1-4) constructed for the confirmation of the key fragment of PmFMO3. The full protein sequence of PmFMO3 was divided into 4 fragments depending the same region between *PmFMO3* and *PmFMO24*. Then each of them was replaced with its homolog fragment of *PmFMO24*. **(f)** The confirmation of the key fragment of PmFMO3 for the hydroxylation activity in *N. benthamiana* leaves. The separative fragments were cloned from the vector of *PmFMO3* and *PmFMO24*. Different mutations were structured by the combinations of different fragment. Then, they were overexpressed in *N. benthamiana* leaves, and fed with resveratrol. PmFMO3 and PmFMO24 was used as the positive and negative control, respectively. **(g)** Three types of transcription factors regulate the promoters of *PmSTS1*, *PmFMO3*, and *UGT72B90* by luciferase assay. GUS: *β*-glucuronidase, used for control, WRKY: PM02g6581, NAC: PM02g10052, and MYB: PM11g55274.

To explore the reason of THSG specifically existed in *P. multiflorus*, 12, 25, and 24 FMO family genes were identified in *Beta vulgaris*, *F. esculentum*, and *F. tataricum* genome, respectively (Table S16-18). Phylogenetic analysis indicated that PmFMO3 is closely related to PmFMO24, GWHPBJBK018339 from *F. esculentum* and GWHPBJBL014999 from *F. tataricum* (Fig. 5(c); Fig. S15). Both *PmFMO24* and *PmFMO3* were transposed duplicated from the same ancestral chromosomal location (*PmFMO13*), which is not expressed in any tissue of *P. multiflorus*. Protein alignment analysis showed that the sequences of these four proteins had high similarity (Fig. S16). All these four genes were transiently over-expressed in *N. benthamiana* leaves. When fed with *trans*-resveratrol, THSG was only remarkably detected in the *PmFMO3*-overexpression leaves, while the *PmFMO24*, *GWHPBJBK018339*, or *GWHPBJBL014999*-overexpression leaves showed no difference with the control leaves (Fig. 5(d)). This is consistent with the result that no THSG can be detected in *F. esculentum*, and *F. tataricum* (Fig. S17). Then we speculated that neofunctionalization may lead to the capacity to biosynthesize THSG in *P. multiflorus*. To determine the key sites of PmFMO3, the protein sequence of PmFMO3 was divided into four parts, and each of them was replaced with the corresponding segment of PmFMO24 to make four different mutations (Fig. 5(e)). All the mutations showed no hydroxylation activity to *trans*-resveratrol (Fig. 5(f)). It seemed that the hydroxylation activity of PmFMO3 to resveratrol was dependent on the overall three-dimensional structure, and segment replacement will destroy its activity.

### 3.6. The transcription factor identification of the stilbenoid gene cluster in *P. multiflorus*

Due to PM02g6581 (WRKY), PM02g10052 (NAC), and PM11g55274 (MYB) were speculated to involve in the regulation of THSG biosynthesis by WGCNA (Fig. 2(d)), we tried to explore whether they regulate the expression of this gene cluster. The 2 kb upstream regions of *PmSTS1*, *UGT72B90*, and *PmFMO3* were cloned and analyzed, MYB, NAC, and WRKY binding domains were predicted to exist in their promoters (Table S19-21). By luciferase assay, PM02g10052 (NAC) were preliminarily verified to positively regulate the expressions of *PmSTS1* and *UGT72B90*, and PM11g55274 (MYB) was preliminarily verified to strongly positively regulate the expressions of *PmSTS1* and *PmFMO3* (Fig. 5(g)).

### 3.7. The substrate selectivity of PmFMO3

Because PmFMO3 was the first stilbenoid-2-hydroxylase, the identification of its substrate selectivity will facilitate the hydroxylated modification of other stilbenoids besides resveratrol. We first selected *cis*-resveratrol (1), pinosylvine (2), dihydroresveratrol (3), oxyresveratrol (4), piceatannol (5), desoxyrhapontigenin (6), pinostilbene (7), pterostilbene (8), piceid (9), and mulberroside A (10) to represent different kinds of stilbenoids (Fig. 6). In *N. benthamiana* leaves, individual PmFMO3 could catalyze the C-2 hydroxylation of 2, 3, 4, 5, and 6, and existed as corresponding glycosides (Figs. S18-22). However, PmFMO3 could not catalyze the hydroxylation of 1, 7, and 8 (Fig. 6). Interesting, when 9 and 10 were used as the substrate in *N. benthamiana* leaves, a small amount of THSG and the catalysate same as of 4 were detected, respectively. While there was no catalysate by *in vitro* enzymatic reaction (Fig. S23). We speculated 9 and 10 may be hydrolyzed by the endogenous glycosidase to release corresponding aglycones (*trans*-resveratrol, and 4), then hydroxylation by PmFMO3 in *N. benthamiana*. To sum up, the hydroxylated and methylated modification of B ring and the hydrogenation of ethylene do not affect the activity of PmFMO3 to hydroxylate the C-2 position of A ring. But when substrate was *cis*-type or A ring was methylated or glycosylated, PmFMO3 could not catalyze them. Furthermore, other types of compounds with the similar structure, such as tyrosine, *p*-coumaric acid, caffeic acid, 5-propylbenzene-1,3-diol, and naringenin, were also be detected as the substrate (Fig. S24). But no corresponding hydroxylation products were detected. PmFMO3 displayed substrate specificity to stilbenoids, which is different from the other stilbenoid-hydroxylase verified in prokaryotes and human.

**Fig. 6.**
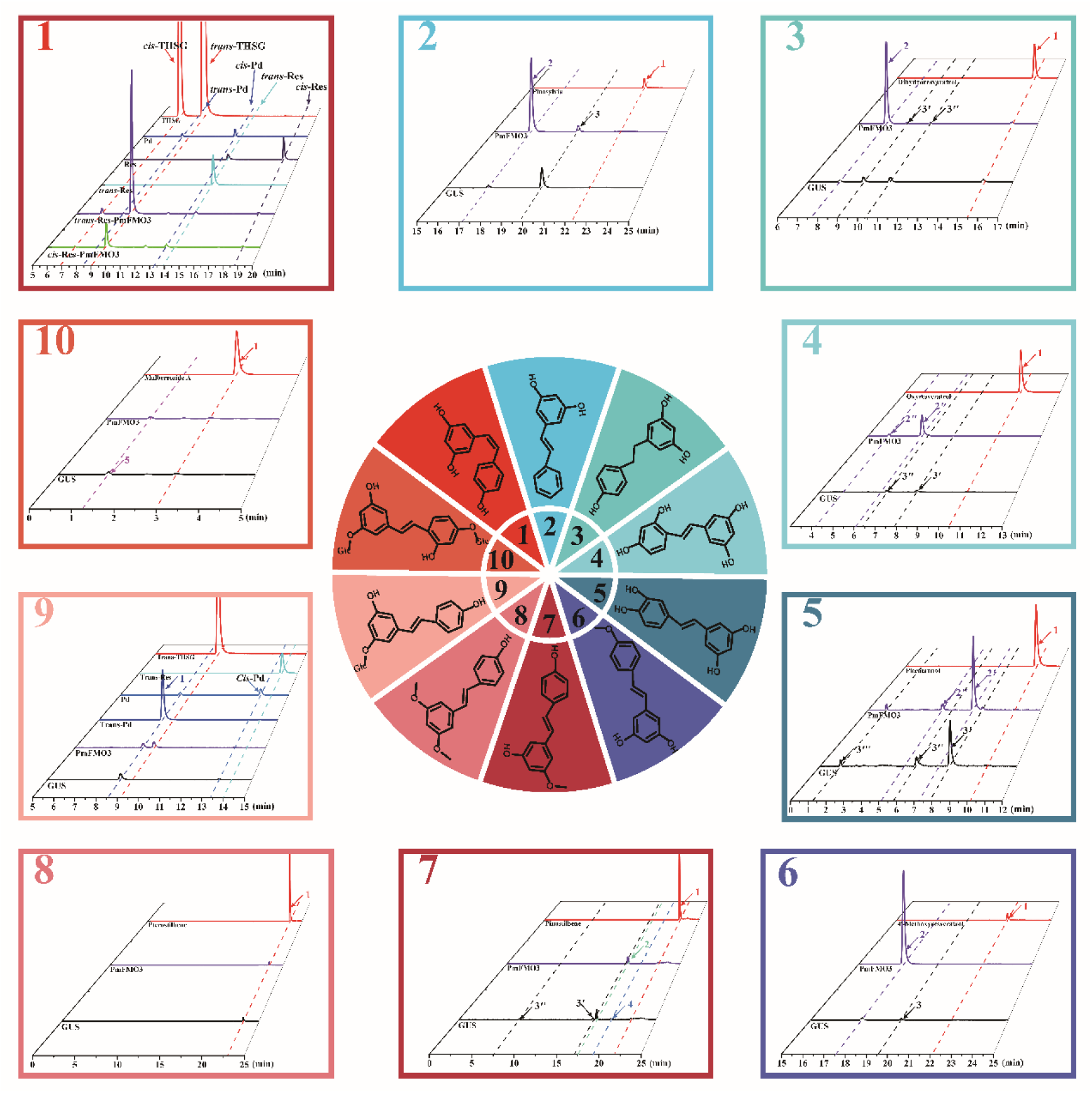
The substrate selectivity of PmFMO3. The blue sector indicates that the stilbenoids can be used as the substrates of PmFMO3, the red sector indicates that the stilbenoids cannot be used as the substrates of PmFMO3. The peak 1 indicates the substrate fed to the *N. benthamiana* leaves; the peak 2 indicates the glycoside of hydroxylation product; the peak 3 indicates the glycoside of the substrate.

## 4. Discussion

Although the dried root and caulis of *P. multiflorus* have been utilized as a traditional medicine for over 1000 years in China, the evolutionary classification of *P. multiflorus* is controversial. Over the centries, it has been successively classified into the *Polygonum*, *Reynoutria*, *Pleuropterus*, and *Fallopia* genus by different plant taxonomists. The *Pleuropterus* genus, which contains *P. multiflorus* and *P. ciliinervis*, was confirmed by combined analysis of chloroplast and nuclear DNA sequence (matK, trnL-trnF, and nrITS) [52]. This classification has been accepted by *China Checklist of Higher Plants* in recent years. The high-quality chromosomal-level reference genome sequence of *P. multiflorus* provided a great opportunity to gain a complete understanding of its evolution. In total, 1.30 Gb genome sequences were assembled with a contig N50 of 117.0 Mb. The assembled quality is better than most published medicinal plants. It was interesting that *P. multiflorus* and *R. hastatulus* were clustered into a clade in phylogenomic analyses, while *F. esculentum*, and *F. tataricum* belong to the other clade (Fig. S3). However, the *Rumex* genus is evolutionarily distant from *Polygonum* L. *s*. *lat*., and the members of *Rumex* genus were often used as the outgroup in phylogenetic analysis of *Polygonum* L. *s*. *lat* based on the previous opinion. The accurate classification and evolution of *Polygonum* L. *s*. *lat*. may need more species to have the whole genome sequenced.

The stilbenoids, a group of plant phenolic metabolites derived from phenylpropanoid biosynthetic pathway, are renowned for their health benefits. A total of 459 natural stilbenoids from 45 plant families and 196 plant species have been identified until 2022 [2]. However, the understanding of biosynthetic pathway of stilbenoids was limited to the synthesis of resveratrol by STS, the decorating enzymes that synthesize the resveratrol derivatives are rarely identified in plants. THSG, the glucoside of C2-hydroxylated resveratrol derivative, is specifically enriched in *P. multiflorus*. Resveratrol had been verified as involved in the THSG biosynthetic pathway by stable isotope labelling [9], and only several STSs were identified and characterized in *P. multiflorus* [46,47,53]. Five types of enzymes have been reported to catalyze aromatic hydroxylation, including CYP450, FMO, nonheme mononuclear iron dioxygenase, pterin-dependent nonheme monooxygenase, and diiron hydroxylase [54,55]. Among them, some prokaryotic and human CYP450 and FMO family members had been reported could specifically hydroxylate the C-3′ position of the B ring of stilbenoids [44,48–51,56]. In the present work, a new type of gene cluster in Chr02 containing an STS (PmSTS1), a FMO (PmFMO3), and a UGT (UGT72B90) were identified to be involved in the THSG biosynthesis (Fig. 4(b)). In recent years, many plant metabolic gene clusters have been identified [16,17]. They are involved in the biosynthesis of terpenoids [18], alkaloids [19], fatty acids [20], benzenoids [21], cyanogenic glycosides [22], flavonoids [23], and phenolamides [24,25]. This is the first stilbenoid gene cluster to be discovered in plants. The genes of this stilbenoid gene cluster were regulated by MYB and NAC type transcription factors (Figs 2(d), 5(g)), and NAC type transcription factor was first found to be involved in the regulation of stilbenoid biosynthesis. The formation mechanisms of plant metabolic gene clusters are not fully understood. Some reports showed TE-mediated chromosome recombination and gene relocation may contribute to cluster formation in eudicots [57–59]. In the present work, many TEs also dispersed in the cluster, especially four Copia-type retrotransposons, which were inserted in the first intron of *PmFMO3* (Fig. 4(b)). And a higher TE/gene density ratio was observed within the location of the gene cluster (Fig. 5(b)). Both *PmFMO3* and *UGT72B90* are transposed duplicates. All these phenomena implied that this stilbenoid gene cluster may also form by TE-mediated gene duplications and neofunctionalization.

FMOs are a class of enzymes that can transfer hydroxyl groups to a large variety of small, nucleophilic, heteroatom-containing substrates [60]. They use flavin as an electronic medium and circulate between the oxidation state and reduction state for electron transfer but not metal [61]. FMOs were discovered during the 1960s in hepatic microsomes as their detoxification of a vast spectrum of xenobiotics [62. FMOs have also been found involving the biosynthesis of specialized metabolism in bacteria and unicellular eukaryotic organisms [55,63,64]. But until recently very little was known about the function of FMOs in plants [65]. Here we reported PmFMO3 can selectively hydroxylate the C-2 position of resveratrol A ring, which is formed by the condensation and cyclization of three malonyl-CoA [7]. The C-3 and C-5 positions of the A ring are also hydroxyl groups. It has previously been shown that hydroxylation of aromatic rings catalyzed by microbial FMO is stimulated by deprotonation of an unmodified hydroxyl group *ortho* or *para* to the target position and the subsequent electrophilic attack by the reactive flavin C4a-hydroperoxide intermediate [66–68]. The C-2 hydroxylation of resveratrol by PmFMO3 is also consistent with the law. Accidentally, when PmFMO3 expressed in prokaryotic expression system was added to the *in vitro* reaction mixture, no catalysate can be detected. While when UGT72B90 was also added to the reaction mixture, THSG can be significantly detected (Fig. 4(d)). We reviewed all the articles related to the isolation and identification of chemical constituents from *P. multiflorus*, and found that the aglycone of THSG have not been detected and isolated from *P. multiflorus*. And the aglycone could not be detected during THSG hydrolysis [69]. So, we speculated that the aglycone of THSG may be unstable, and should be stabilized by glycosylation. This speculation was also confirmed by the result of computational chemistry (Fig. S13).

The position and number of hydroxyl substituent of stilbenoid affect drug potency and efficacy, so many experts tried to site-specifically increase the hydroxyl substituent number of stilbenoids by chemical synthesis or biological synthesis, but the modification sites most focused on the 3, 4, and 5 position of A ring or B ring [70,71]. As the first stilbenoid hydroxylase derived from plants and the modification site was C-2 position of A ring, we tested the substrate selectivity of PmFMO3. By using different kinds of stilbenoids as the substrates, we found that the hydroxylated and methylated modification of B ring, which derived from *p*-coumaroyl-CoA and the hydrogenation of ethylene does not affect the hydroxylation activity of PmFMO3, but when the substrate was *cis*-type or A ring was methylated or glycosylated, PmFMO3 could not catalyze them (Fig. 6). Its substrate selectivity to stilbenoids is partly consistent with Sam5, a FMO protein from actinomycetes which could hydroxylate the C-3 position of stilbenoid B ring [44]. Their hydroxylation activity to the target ring of stilbenoids were not influenced by the modification of the other ring. We speculated that the modification of the other ring may do not influence the interaction between the active site residues of FMO protein and the catalytic site of the stilbenoids. But PmFMO3 cannot hydroxylate the other types of structurally similar compound that can be used as substrate of Sam5 [56] and HpaB [72,73], such as tyrosine, *p*-coumaric acid, caffeic acid, and naringenin. We also tested whether PmFMO3 can use 5-propylbenzene-1,3-diol, which the B ring of compound 3 was replaced as the methyl, as the substrate. But the result was negative. Taking into consideration the different origin of A ring and B ring of stilbenoids, we speculate PmFMO3 specifically recognizes the A ring of stilbene skeleton, which formed by condensation and cyclization of three malonyl-CoA.

## Acknowledgements

We thank Professor David R Nelson from the University of Tennessee Health Science Center, and Professor Michael H. Court from Washington State University for CYP450 and UGT family gene naming. We thank Professor Yaqiong Su from Xi’an Jiaotong University for the computational chemistry about the aglycone of THSG. We also thank Mr. Baoying Sun from the Institute of Botany, Chinese Academy of Sciences for the botanical scientific illustration shown as Fig. 2(a). We also thank Dr. Xinyu Zhang from John Innes Centre for the helpful suggestions and language polishing on this manuscript. This work was supported by the National Natural Science Foundation of China (Grant Nos. 32202415), the Scientific and Technological Innovation Project of China Academy of Chinese Medical Sciences (CI2021A04404), and the Fundamental Research Funds for the Central public welfare research institutes (ZZ14-YQ029, ZXKT22002).

## Conflicts of interest

The authors declare no conflict of interest.

## Author contributions

Z.L., and A.L. led the project planning. J.J., and X.L. carried out the experiments. Y.W. performed the bioinformatic analysis. J.J., and X.L. interpreted the data and participated in discussion. J.J. wrote the paper. Z.L., and A.L. revised the paper. All authors read and approved the final manuscript.

## Data availability

All data are available in the manuscript or the Supplementary Information. All sequence reads were deposited in the Chinese National Genomics Data Center database (https://bigd.big.ac.cn/) under BioProject accession number PRJCA026400.

## Notes

### Competing Interest Statement

The authors have declared no competing interest.

