## Supplementary material for "Genome-enabled stilbenoid gene cluster discovery in *Pleuropterus multiflorus*": Fig. S: Supplementary Figure+Method-line number.docx

**Supplementary Figures**

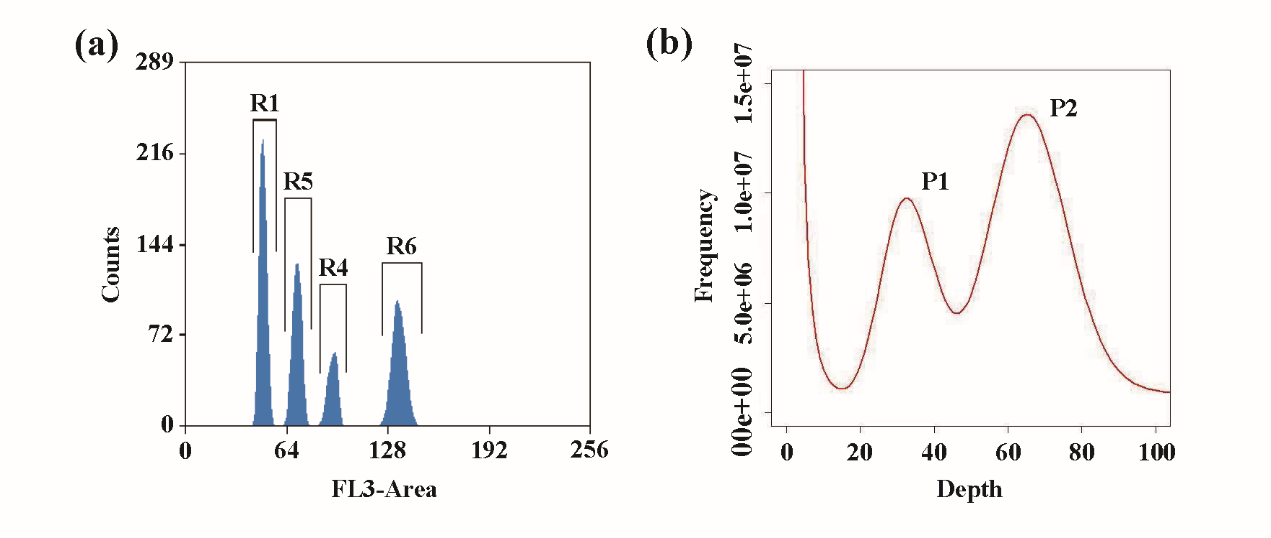

**Fig. S1. Evaluation of the genome size of *Pleuropterus multiflorus* by flow cytometry and K-mer frequency distribution.** **(a)** Fluorescence histograms for genome size evaluation of *P. multiflorus* by flow cytometry. *Solanum lycopersicum* (1C = *c*. 900M), was used as internal reference standard. The fluorescence peaks from G1 and G2-phase nuclei of *S. lycopersicum* were marked as R1 and R4, while the fluorescence peaks from G1 and G2-phase nuclei of *P. multiflorus* were marked as R5 and R6 in pooling samples. **(b)** The K-mer frequency distribution of *P. multiflorus* (K-mer = 19). The X-axis shows the depth of each K-mer, and the Y-axis shows the frequency of each K-mer. P1 represents the hetero-peak, and P2 represents the homo-peak.

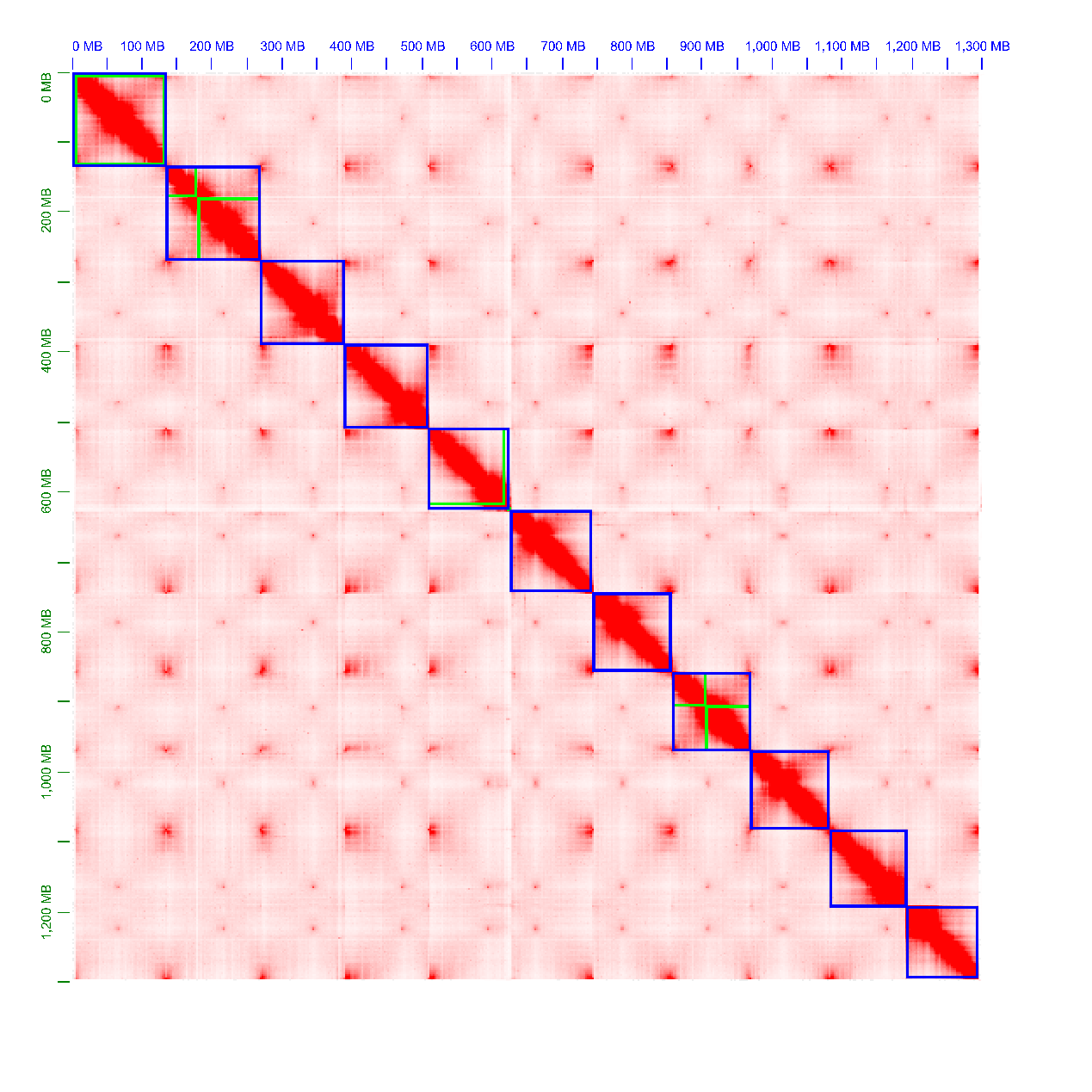

**Fig. S2. Chromosome-level assembly of *Pleuropterus multiflorus* genome using Hi-C technology.**

**Fig. S3. Phylogenetic tree of *P. multiflorus* along with 11 other plants.** Gene family expansions are indicated in brown, and gene family contractions are indicated in blue. The estimated divergence time (million years ago, Mya) is indicated at each node.
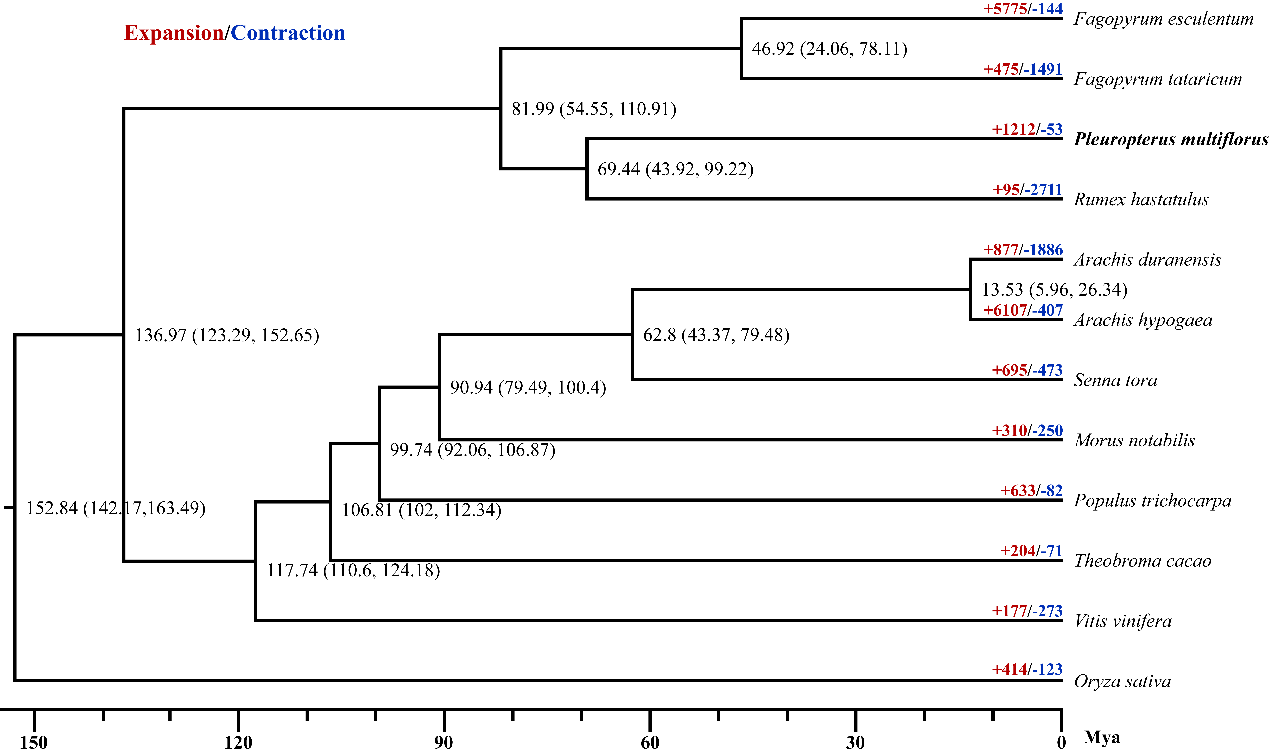

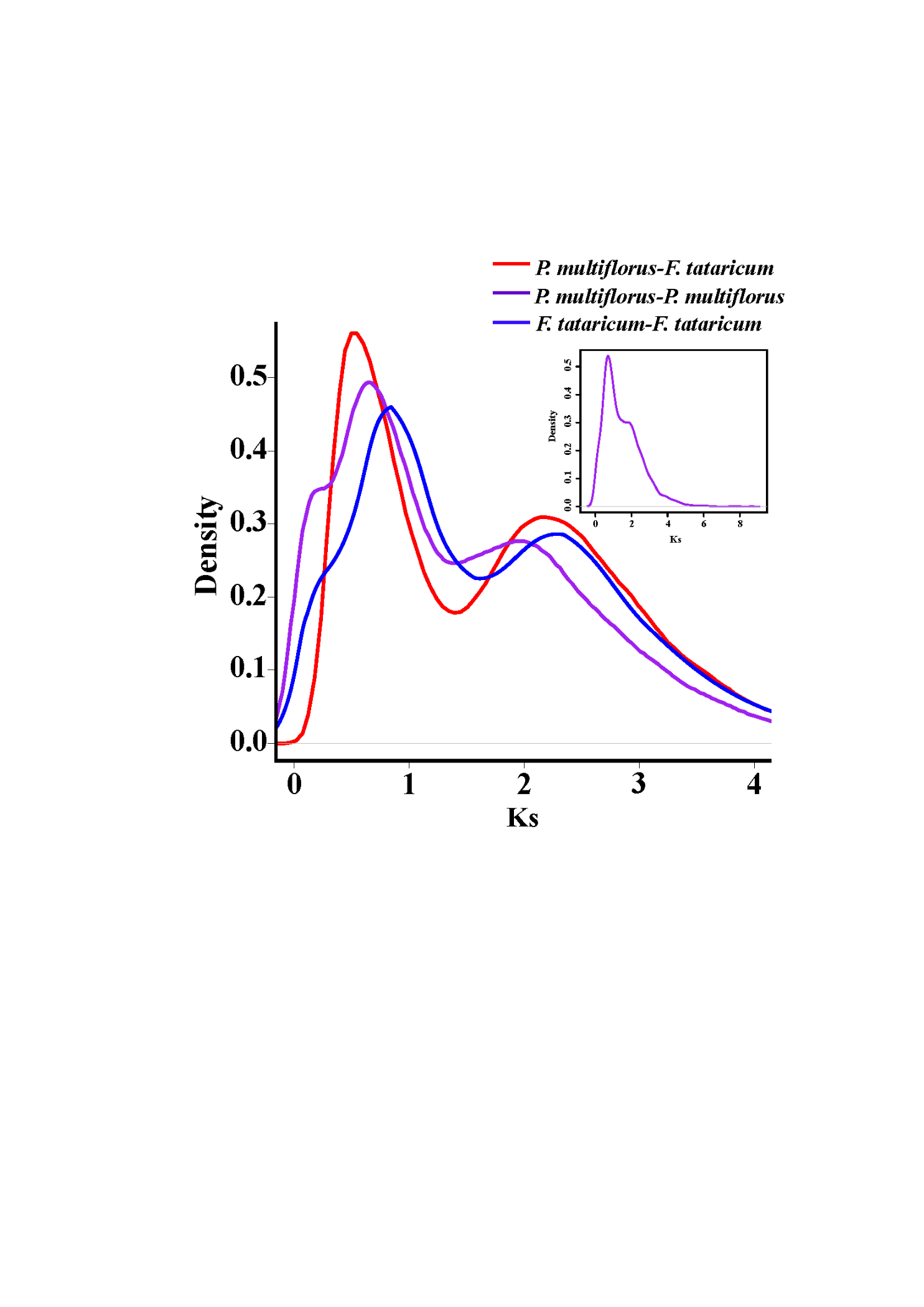
**Fig. S4. The distribution of synonymous substitution rates per gene (Ks) between collinear paralogous genes in *P. multiflorus* and *F. tataricum*.**

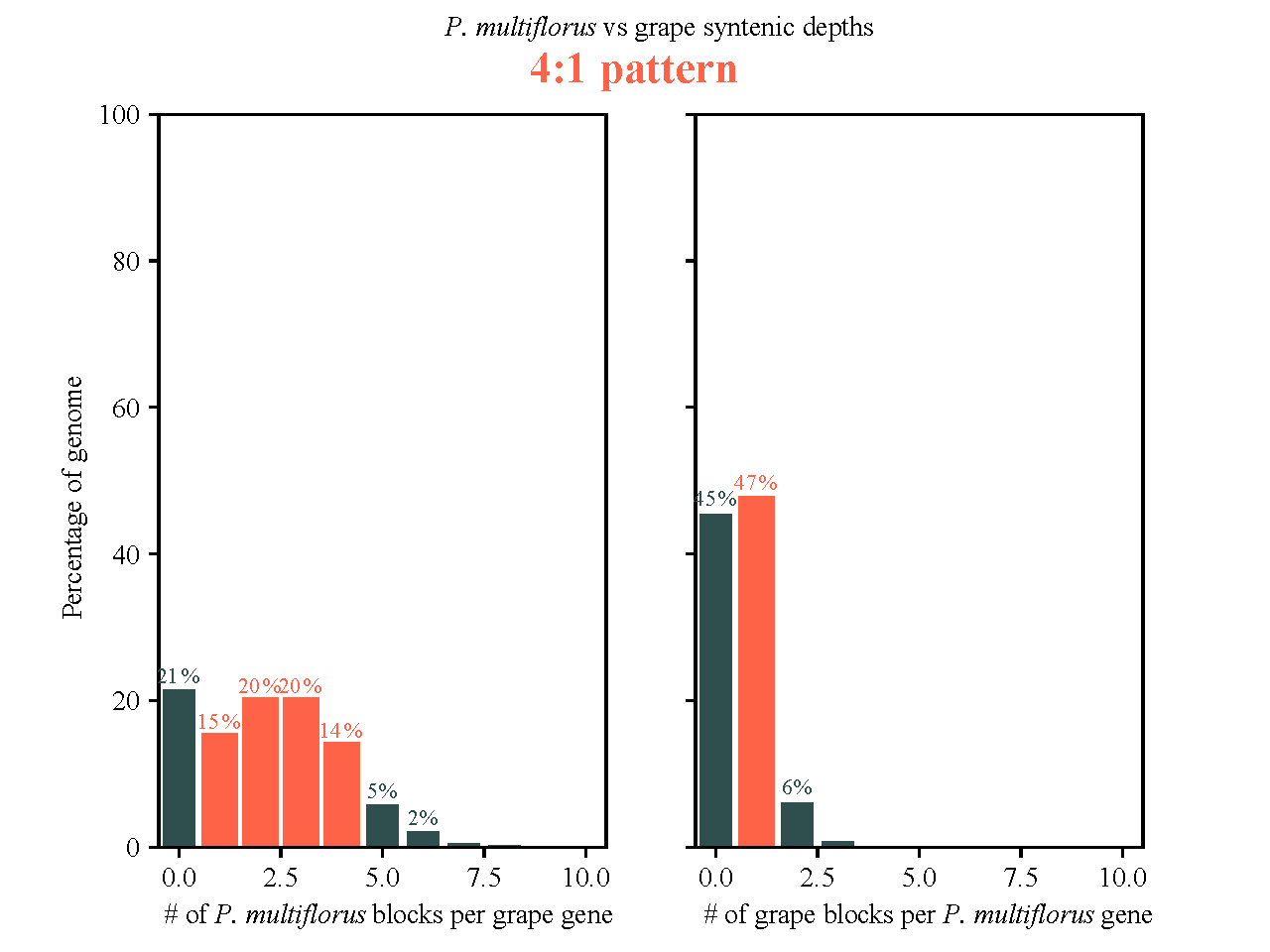
**Fig. S5. Summary of the syntenic analysis between *Pleuropterus multiflorus* and grape.**

**Fig. S6. Summary of the syntenic analysis between *Pleuropterus multiflorus* and pear.**
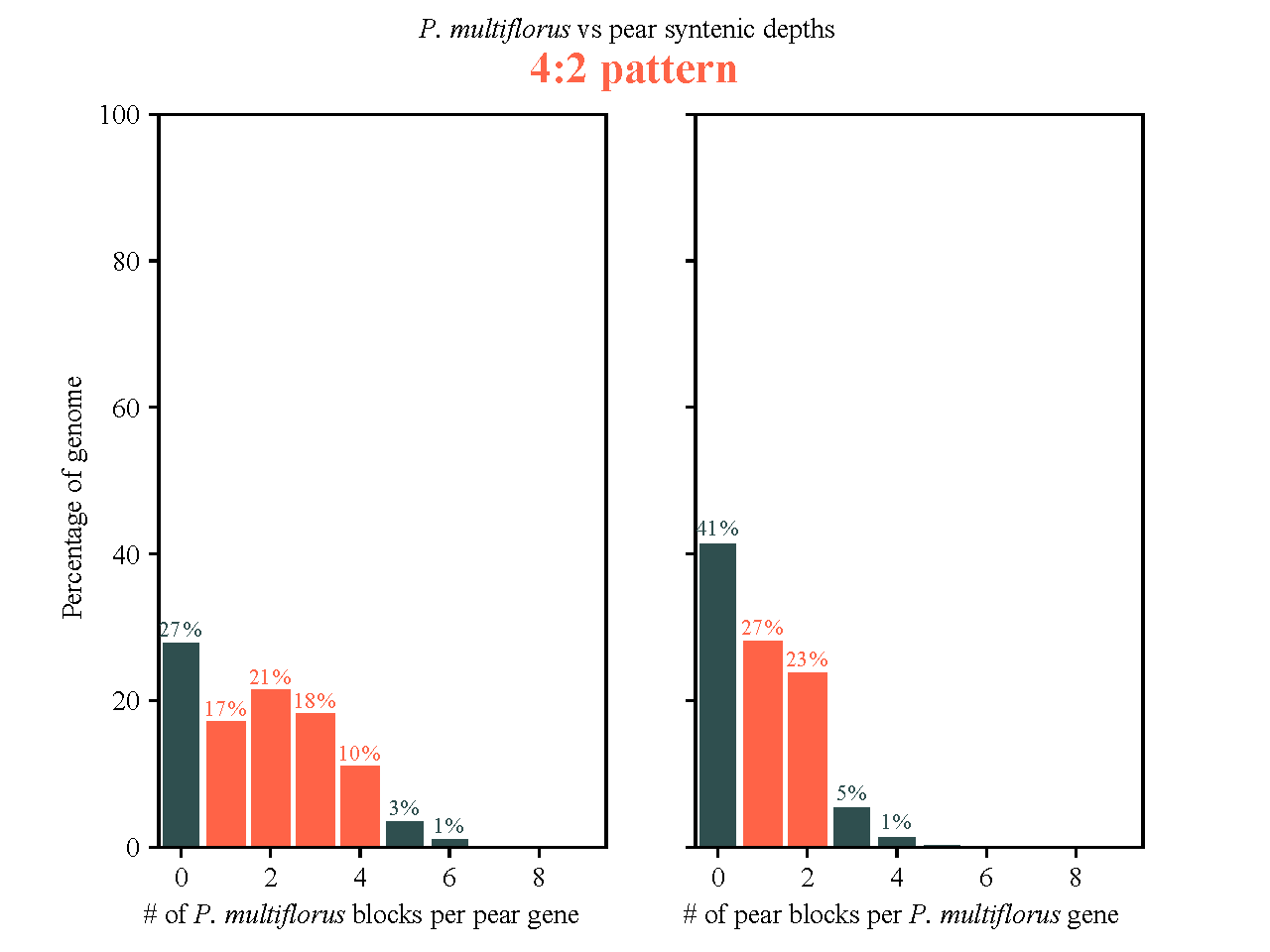

**Fig. S7. Summary of the syntenic analysis between *Pleuropterus multiflorus* and *Fagopyrum tataricum*.**
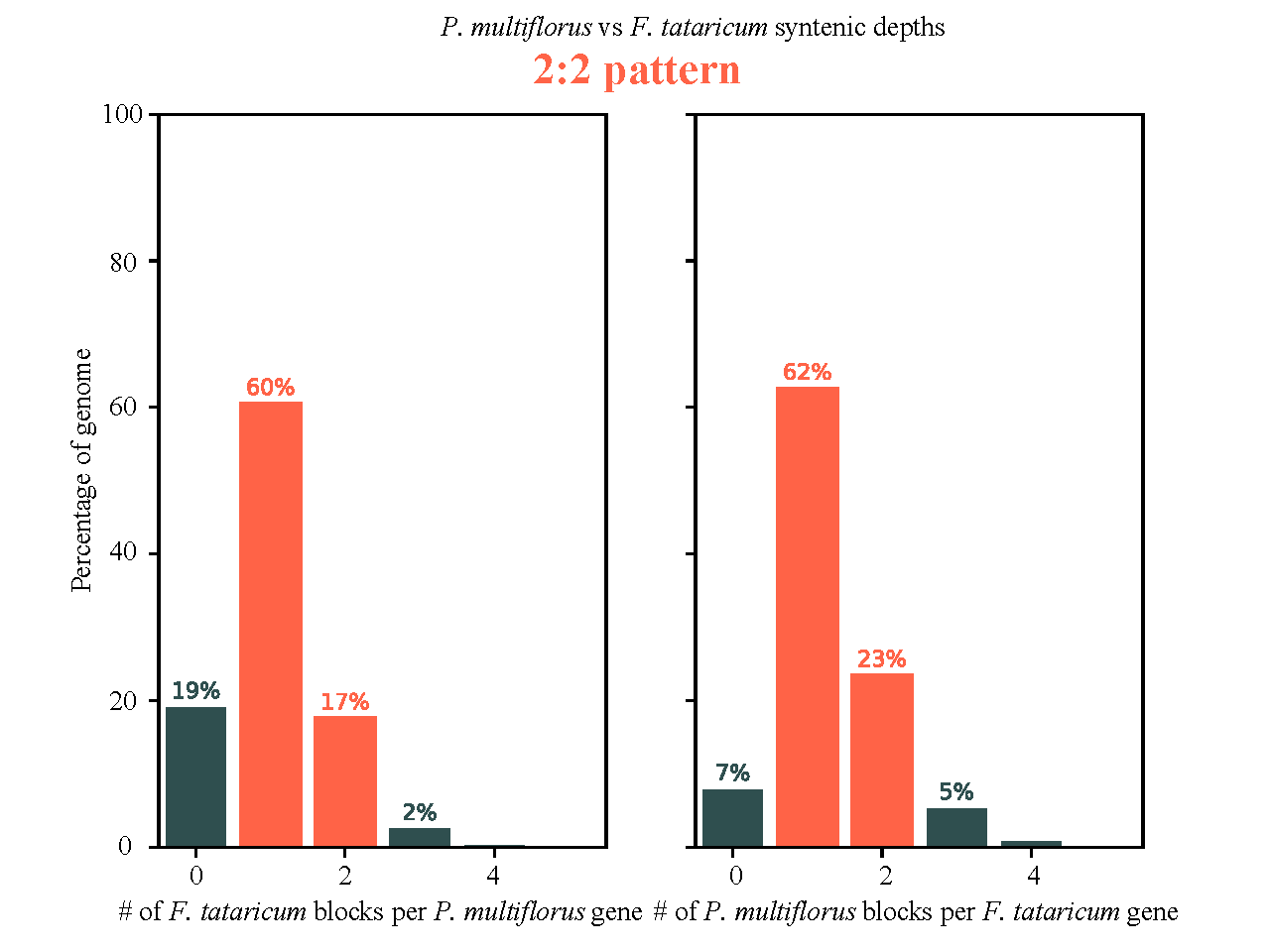

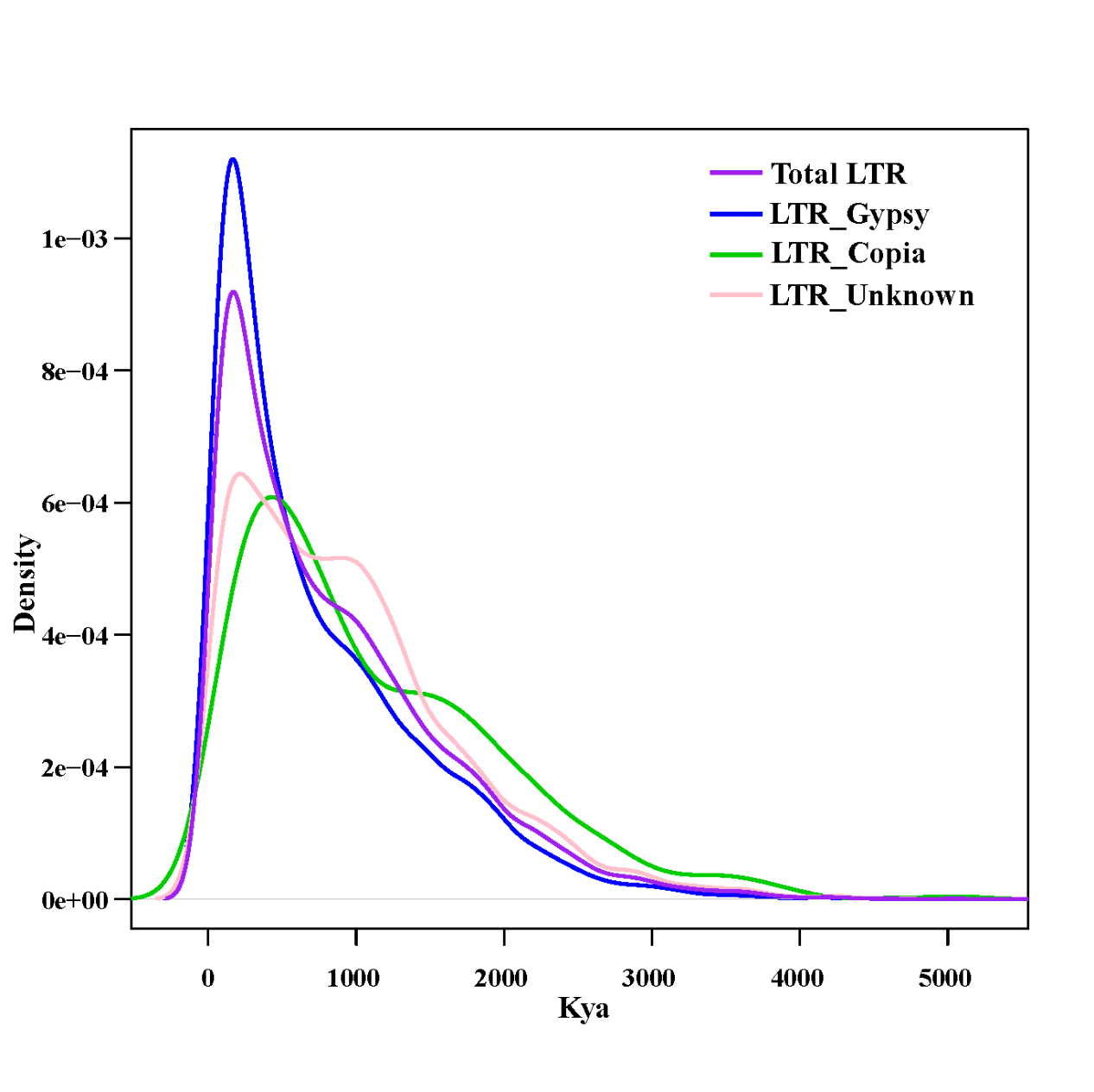
**Fig. S8. Temporal patterns of different type LTR‐RT insertional bursts in *P. multiflorus*.**

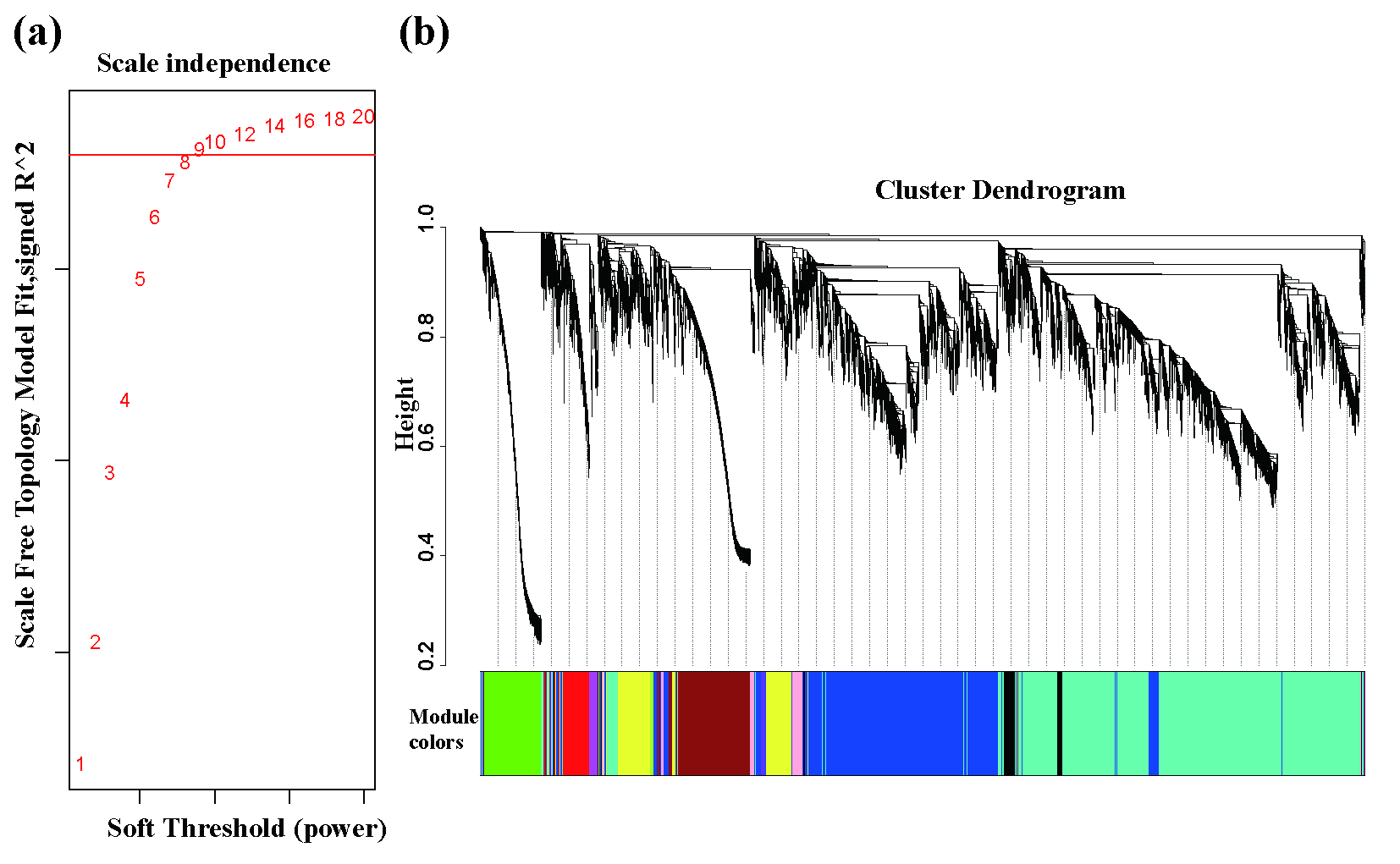

**Fig. S9. Weighted correlation network analysis (WGCNA) for finding the highly correlated genes in *Pleuropterus multiflorus*. (a)** Using the pick Soft Threshold function, a soft threshold of 8 was determined to best fit the network structure. (**b)** Cluster dendrogram constructed by WGCNA show the clustering of expressed genes in 15 *Pleuropterus multiflorus* tissues.

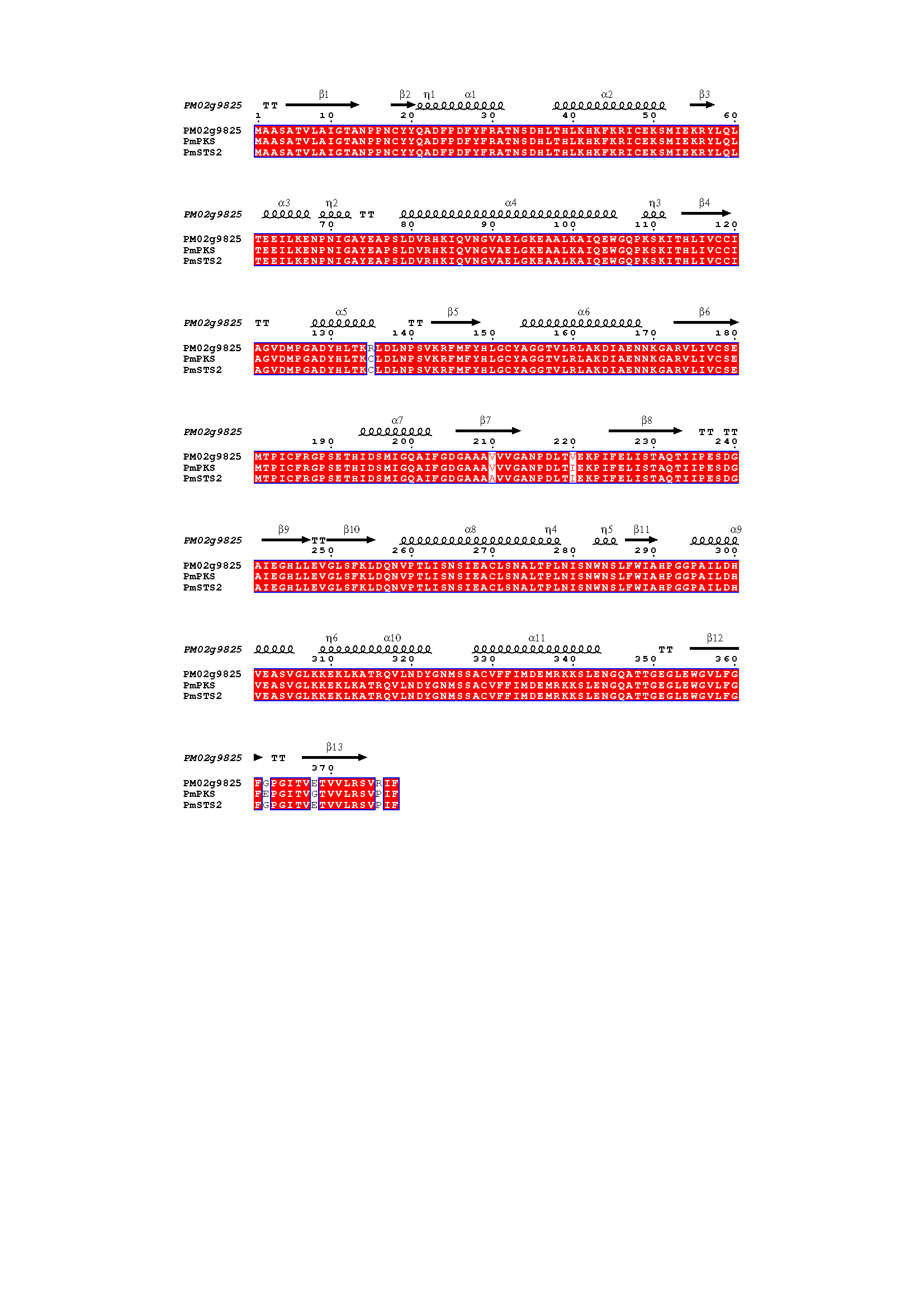

**Fig. S10. Protein similarities between PM02g9825, reported PmPKS, and PmSTS2.** The 3-D structure of PM02g9825 was predicted by Robetta (https://robetta.bakerlab.org/). The alignment was carried out by ClustalW algorithm and visualized by ESPript 3 (https://espript.ibcp.fr/ESPript)

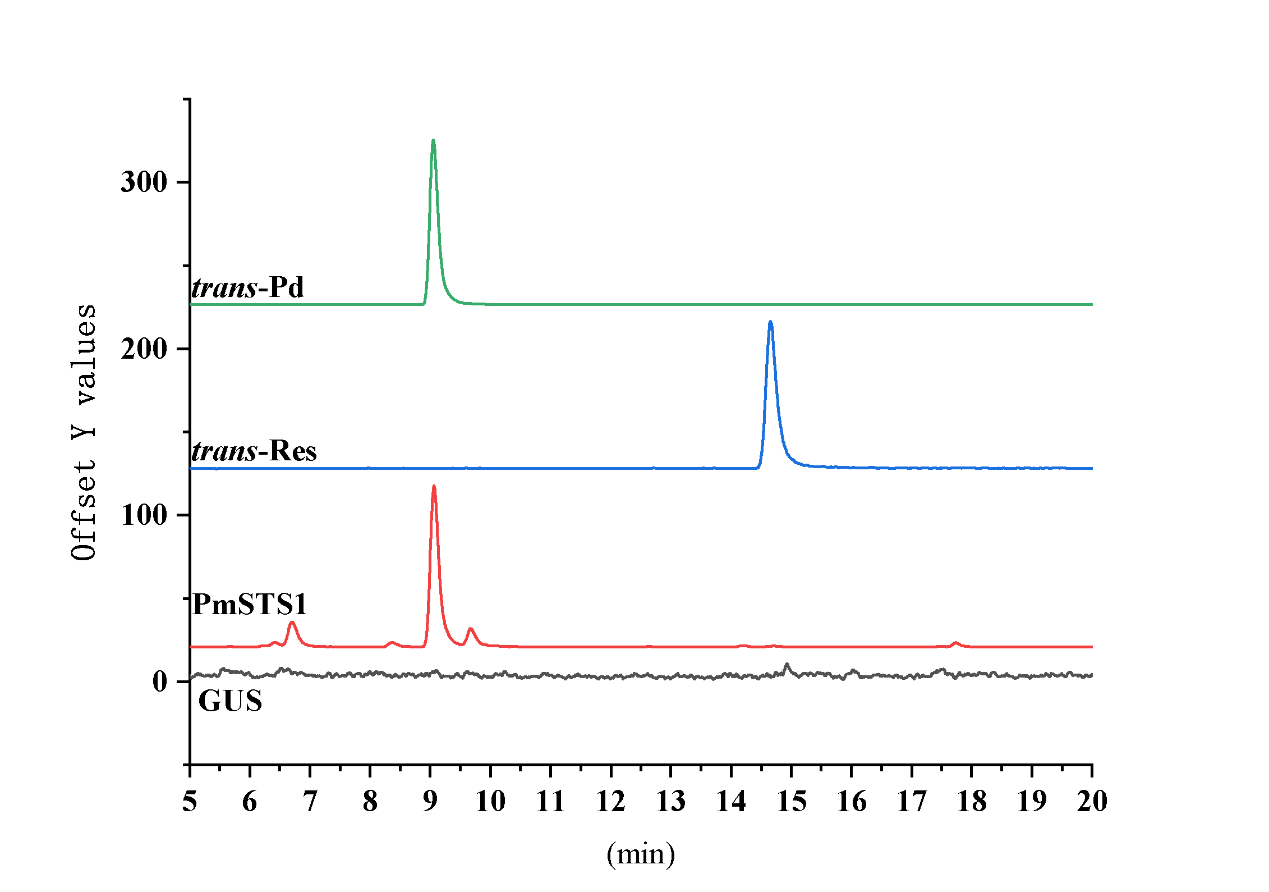
**Fig. S11. Functional verification of PmSTS1 (PM02g9825) in *Nicotiana benthamiana* by UPLC-QQQ-MS.** The black line labeled as GUS represented the stilbenoid type and contents of the control tobacco leaves which overexpressed the GUS protein. The red line labeled as PmSTS1 represented the stilbenoid type and contents of the tobacco leaves which overexpressed *PmSTS1*. The blue and green lines labeled as *trans*-Res and *trans*-Pd represented the elution time of *trans*-resveratrol and *trans*-piceid standards, respectively.

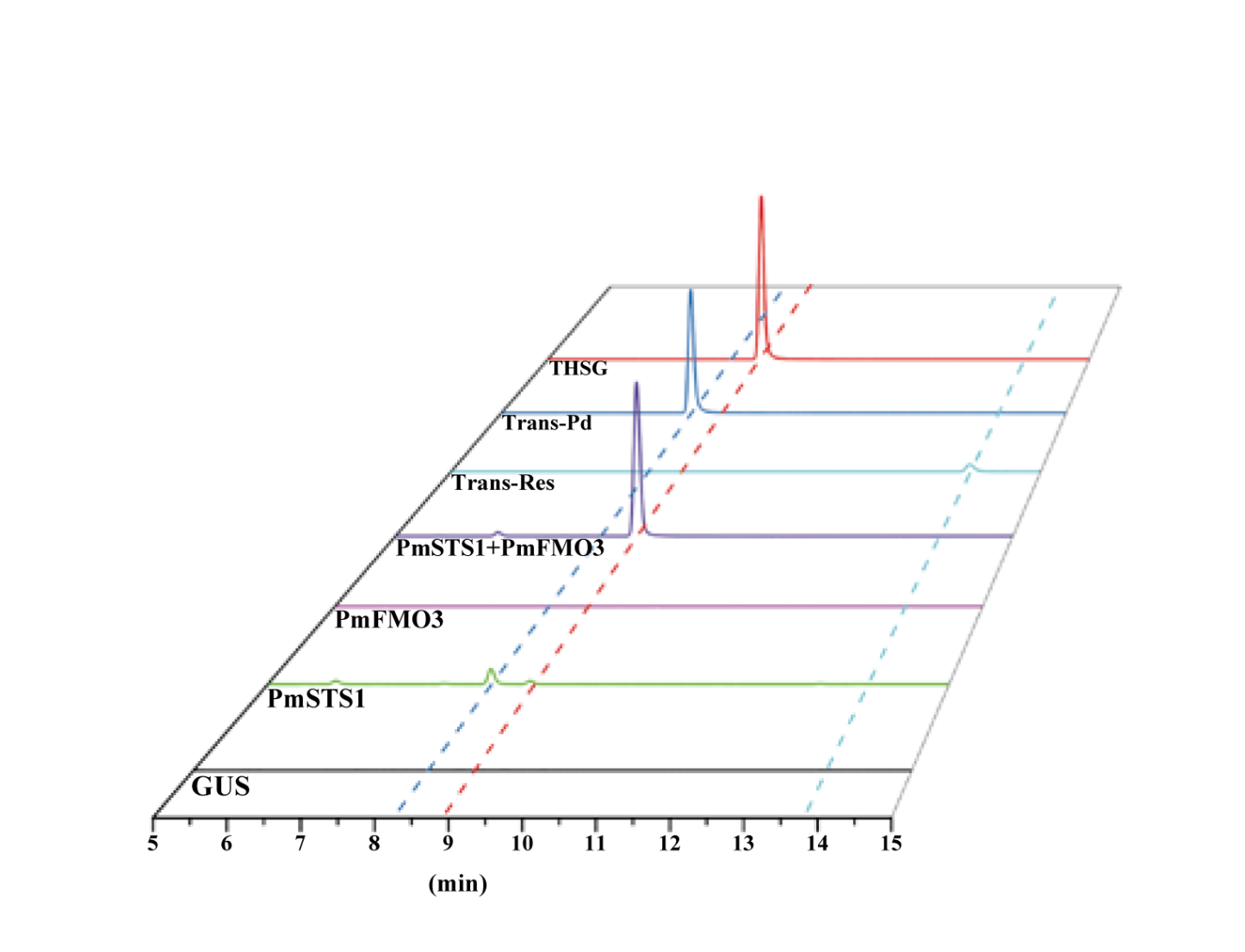

**Fig. S12. The function characterization of PmFMO3 and PmSTS1 separation or combination** **in *Nicotiana benthamiana* by UPLC-QQQ-MS.**

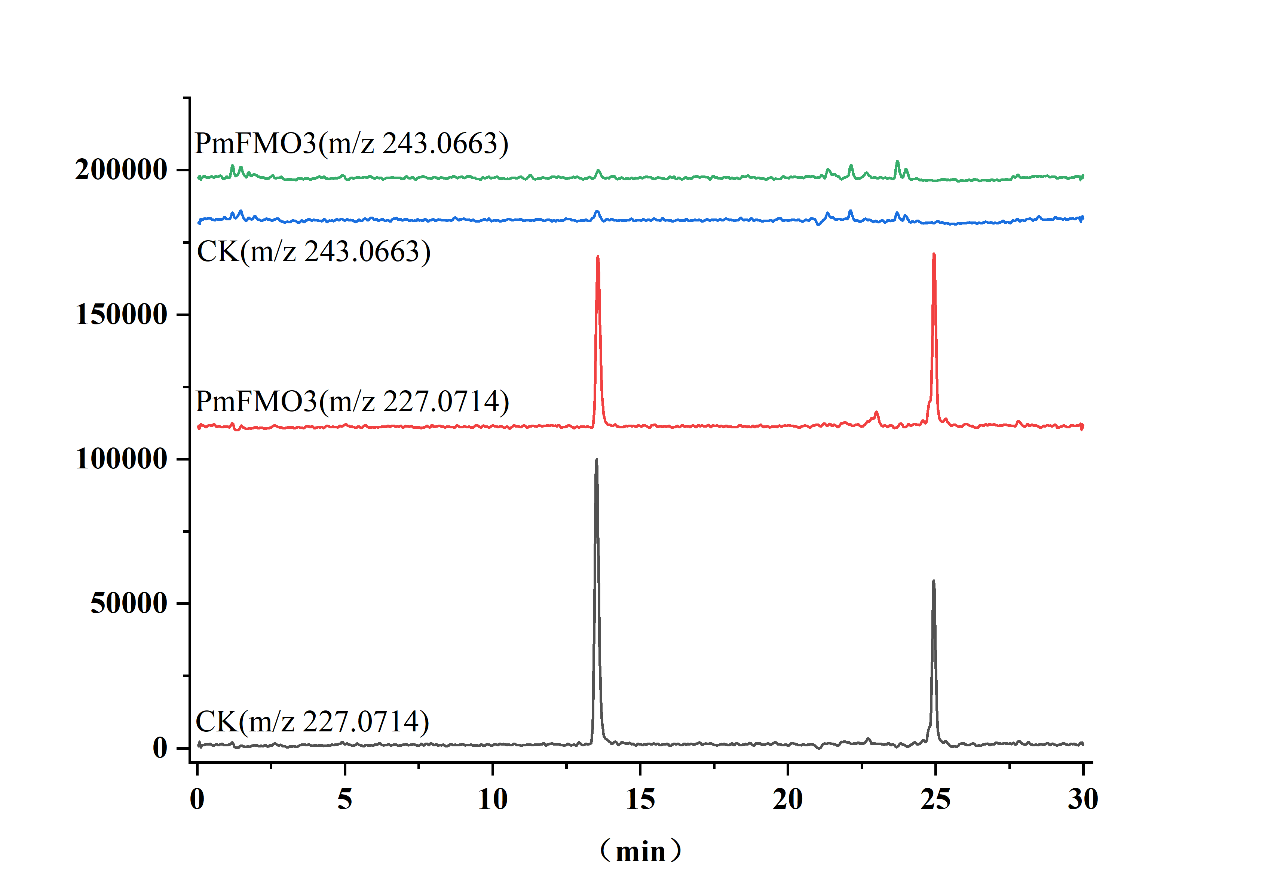

**Fig. S13. Functional verification of PmFMO3 by** ***in vitro* enzyme assay.** The reaction products were detected by UPLC-Q-TOF-MS. The black line labeled as CK (m/z 227.0714) represented the reactive substrate resveratrol added to the reaction system containing the heterologously expressed control enzyme. The red line labeled as PmFMO3 (m/z 227.0714) represented the reactive substrate resveratrol added to the reaction system containing the heterologously expressed PmFMO3 enzyme. The blue line labeled as CK (m/z 243.0663) represented the target traction product THS in the reaction system containing the heterologously expressed control enzyme. The red line labeled as PmFMO3 (m/z 243.0663) represented the target reactive product in the reaction system containing the heterologously expressed PmFMO3 enzyme.

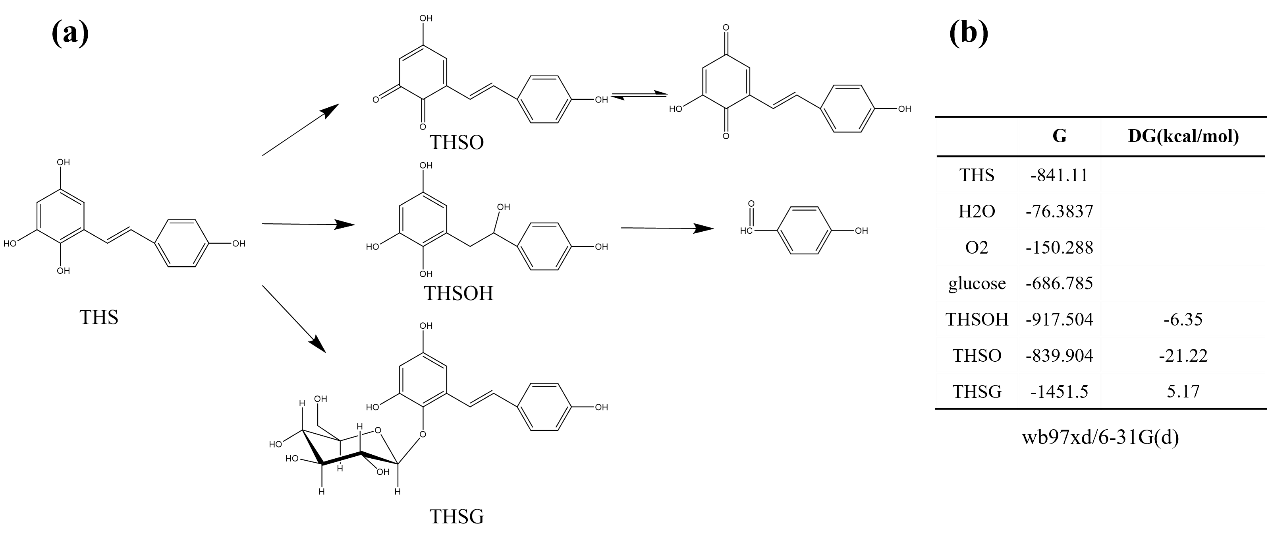

**Fig. S14. Rapid** **transformation and degradation of THS** **and stabilization of THSG validated by computational chemistry.** **(a)** Putative transformation and degradation pathway of THS and stabilization of THSG. **(b)** The Gibbs free energies of different compounds involving the pathway shown in **(a)** were listed.

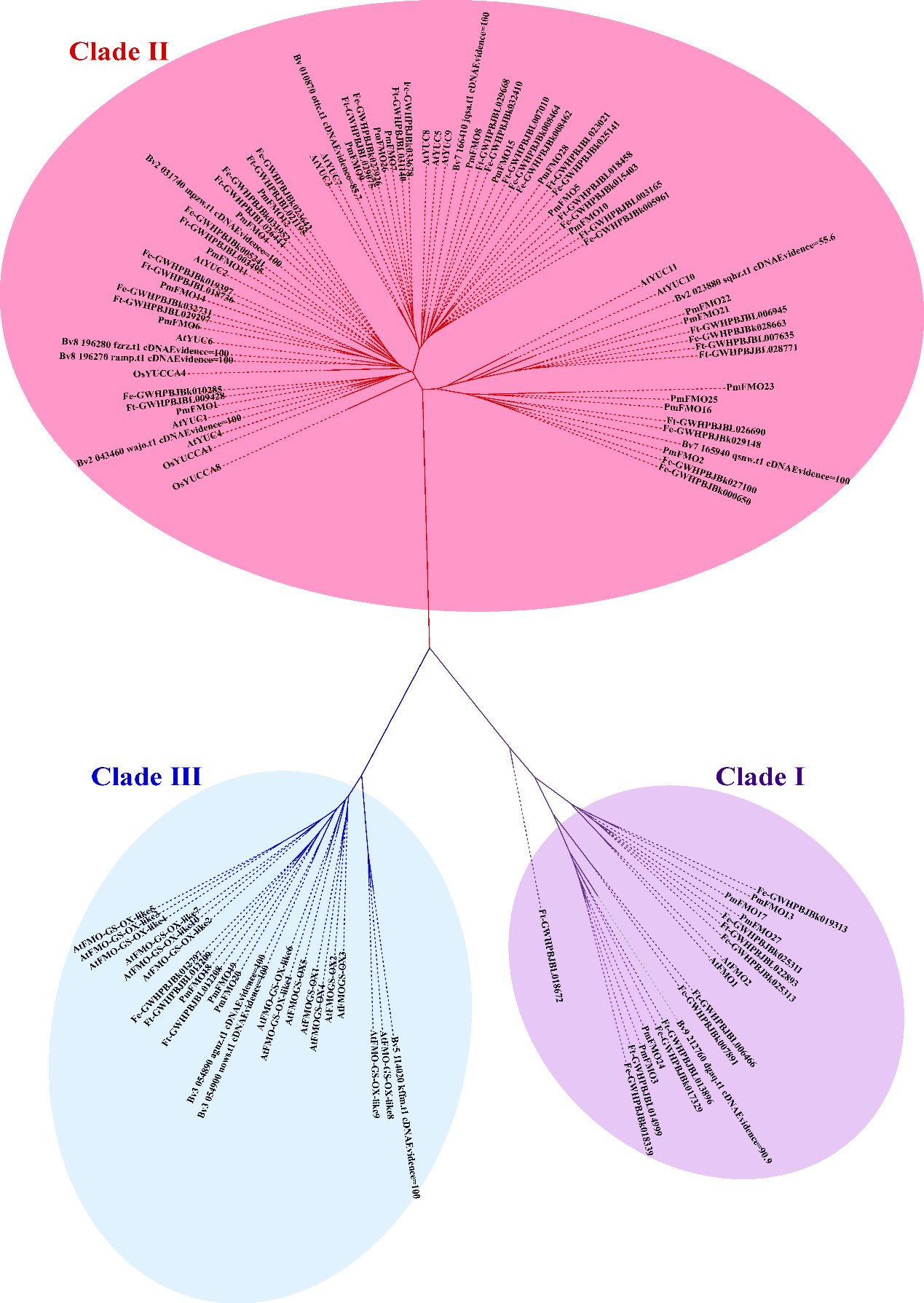
**Fig. S15. Phylogenetic tree of FMO gene families in *P. multiflorus*,** ***Beta vulgaris*,** ***Fagopyrum esculentum*,** ***Fagopyrum tataricum*, *Arabidopsis thaliana*, and *Oryza sativa*.** The phylogenetic tree was built by MEGA11 with NJ method. Different color represented different clade of FMO gene family.

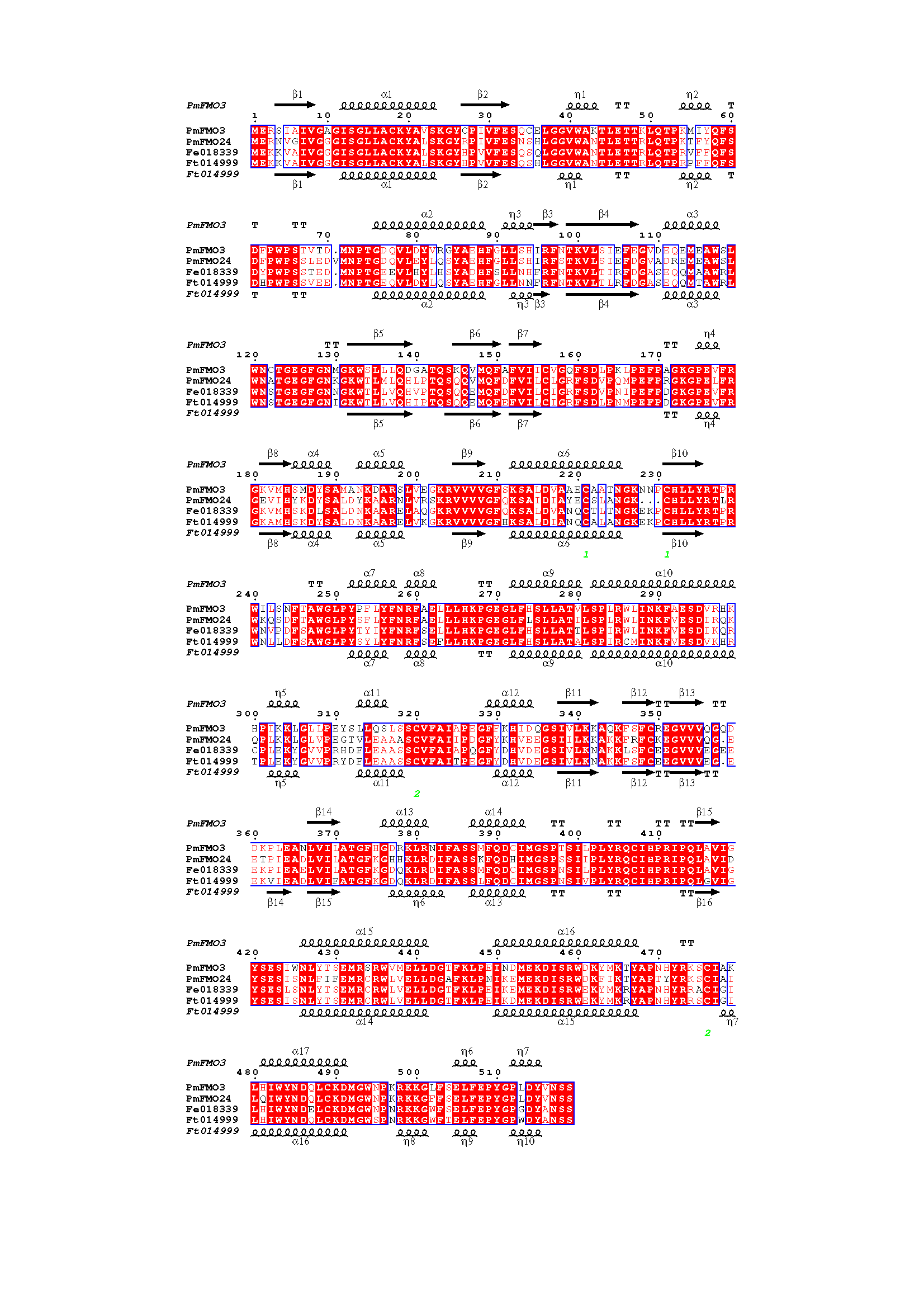

**Fig. S16. Protein similarities between PmFMO3, PmFMO24, GWHPBJBK018339, and GWHPBJBL014999.** The 3-D structure of PmFMO3 and GWHPBJBL014999 (label as Ft014999) was predicted by Robetta (https://robetta.bakerlab.org/). GWHPBJBK018339 was labeled as Fe010339. The alignment was carried out by ClustalW algorithm, algorithm and visualized by ESPript 3 (https://espript.ibcp.fr/ESPript/cgi-bin/ESPript.cgi)

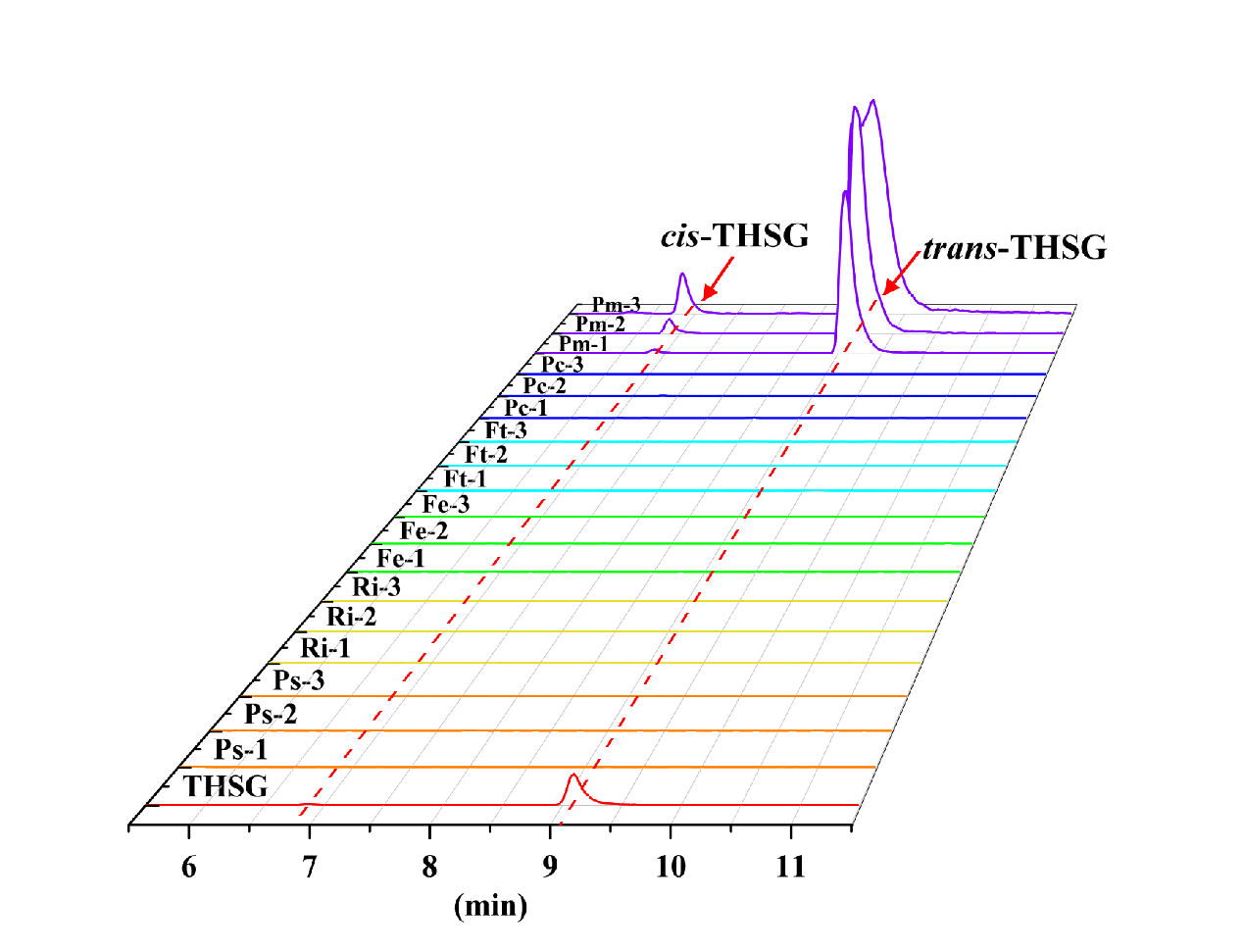

**Fig. S17. The detection of THSG existed in different plant species by UPLC-QQQ-MS.** Some medicinal plant species that had been reported to contain stilbenoids were choosen to detect the THSG. The extraction and detection were performed as the methods shown in the main text. THSG: 2,3,5,4′-tetrahydroxystilbene-2-*O*-*β*-D-glucoside, Ps: *Paeonia* × *suffruticosa*, Ri: *Rubus idaeus*, Fe: *Fagopyrum esculentum*, Ft: *Fagopyrum tataricum*, Pc: *Polygonum cuspidatum*, and Pm: *Pleuropterus multiflorus.*

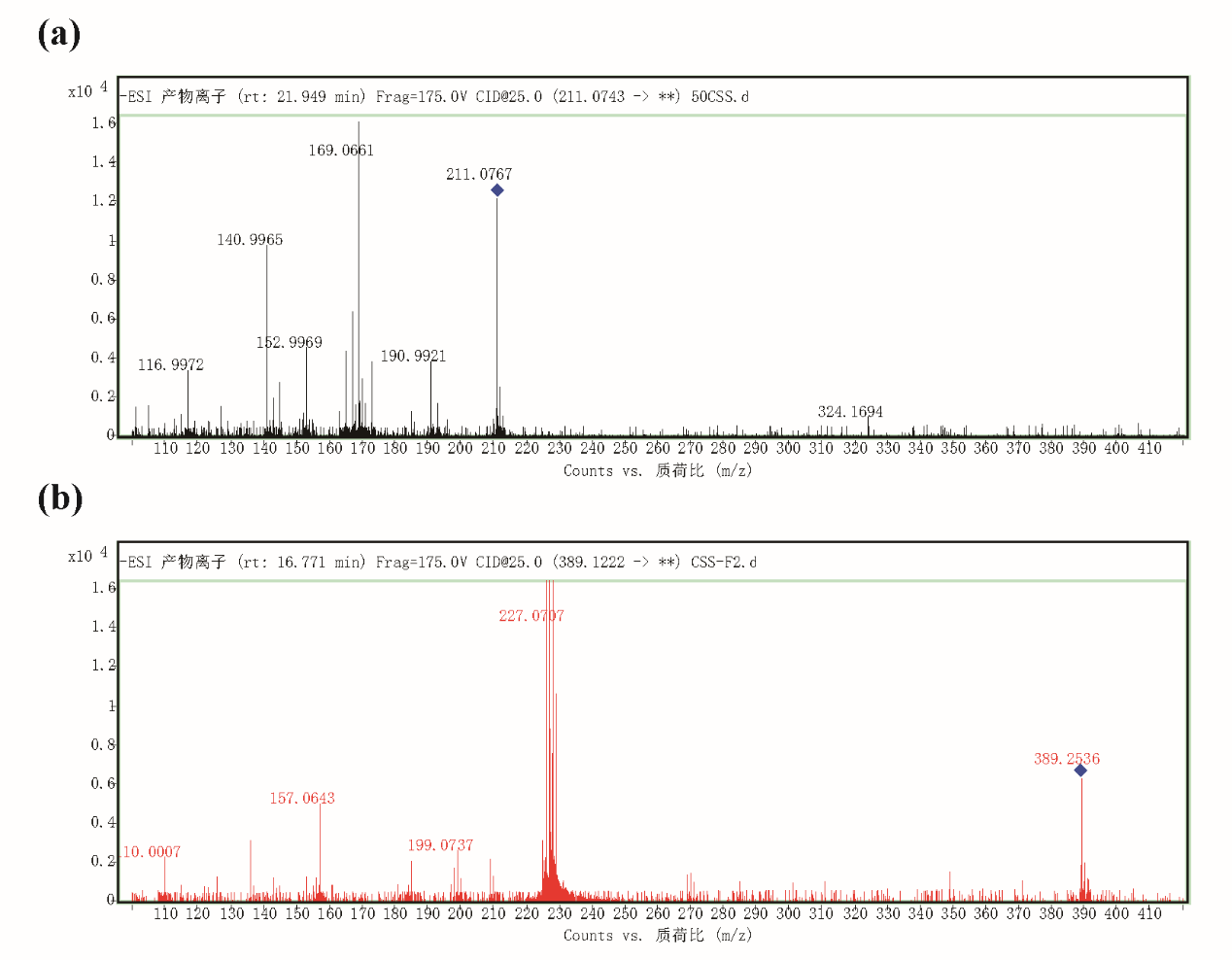
**Fig. S18. The ESI mass/mass spectra of pinosylvine (a) and its hydroxylated glycosylated products (b).**

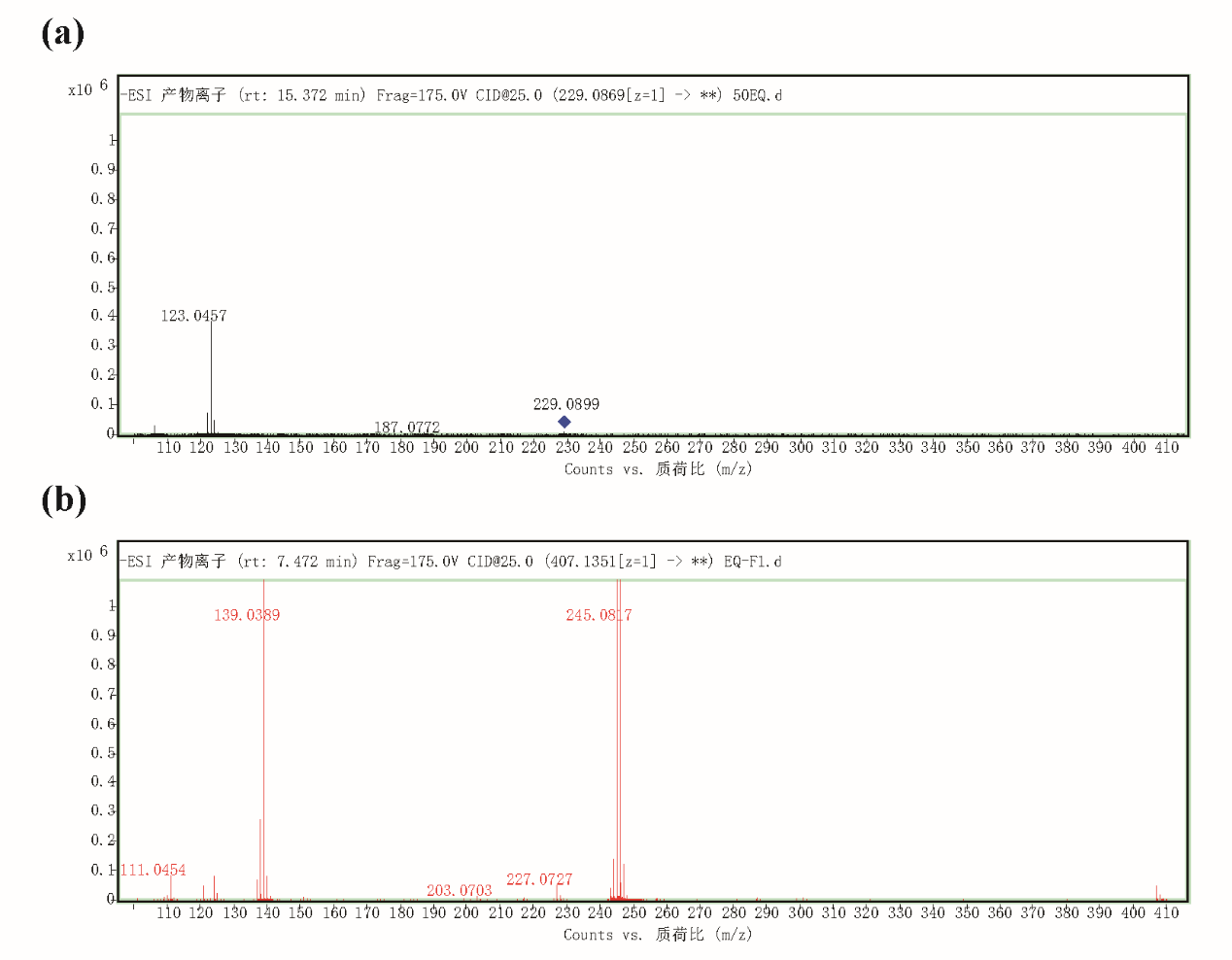

**Fig. S19. The ESI mass/mass spectra of dihydroresveratrol (a) and its hydroxylated glycosylated products (b).**

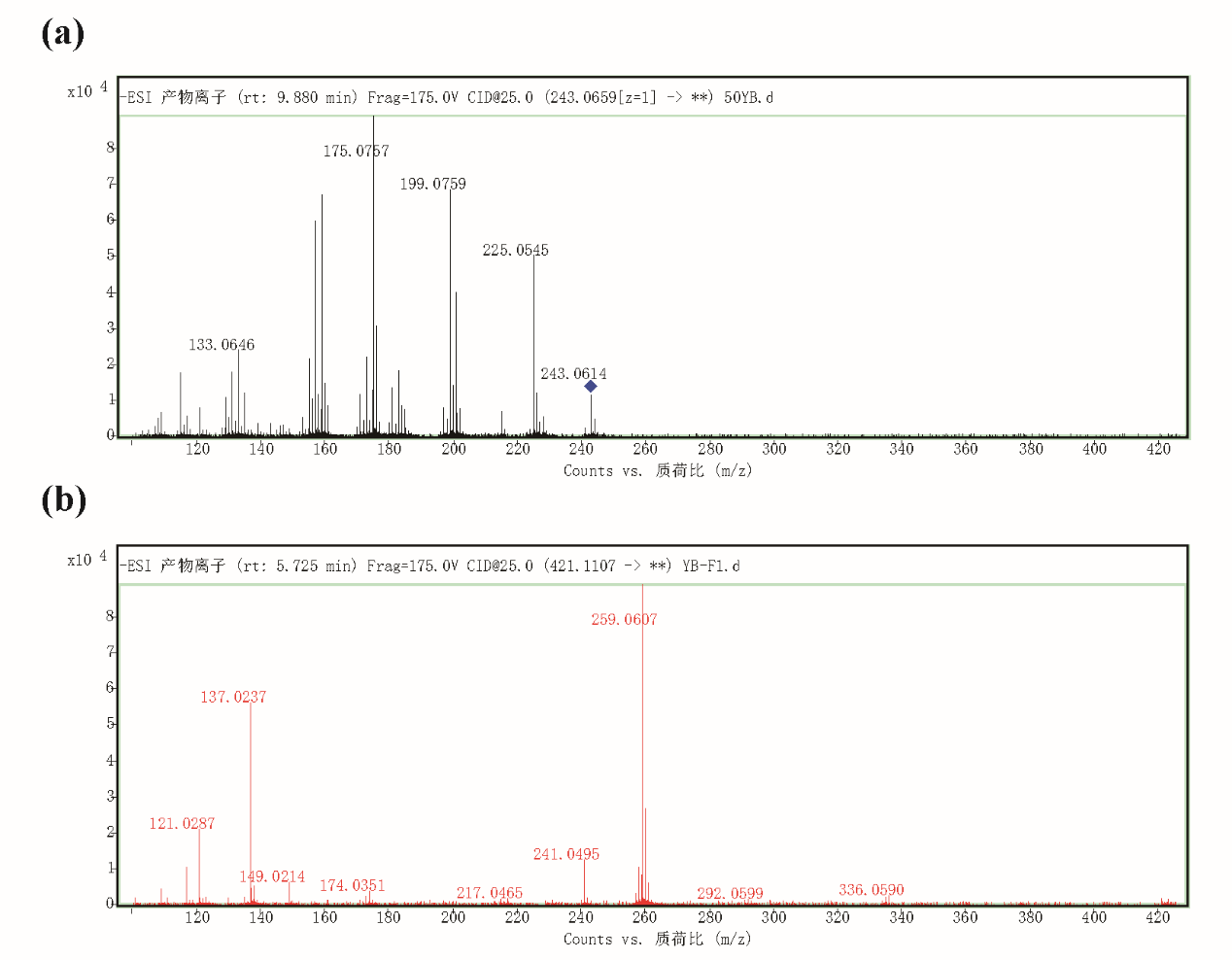

**Fig. S20. The ESI mass/mass spectra of oxyresveratrol (a) and its hydroxylated glycosylated products (b).**

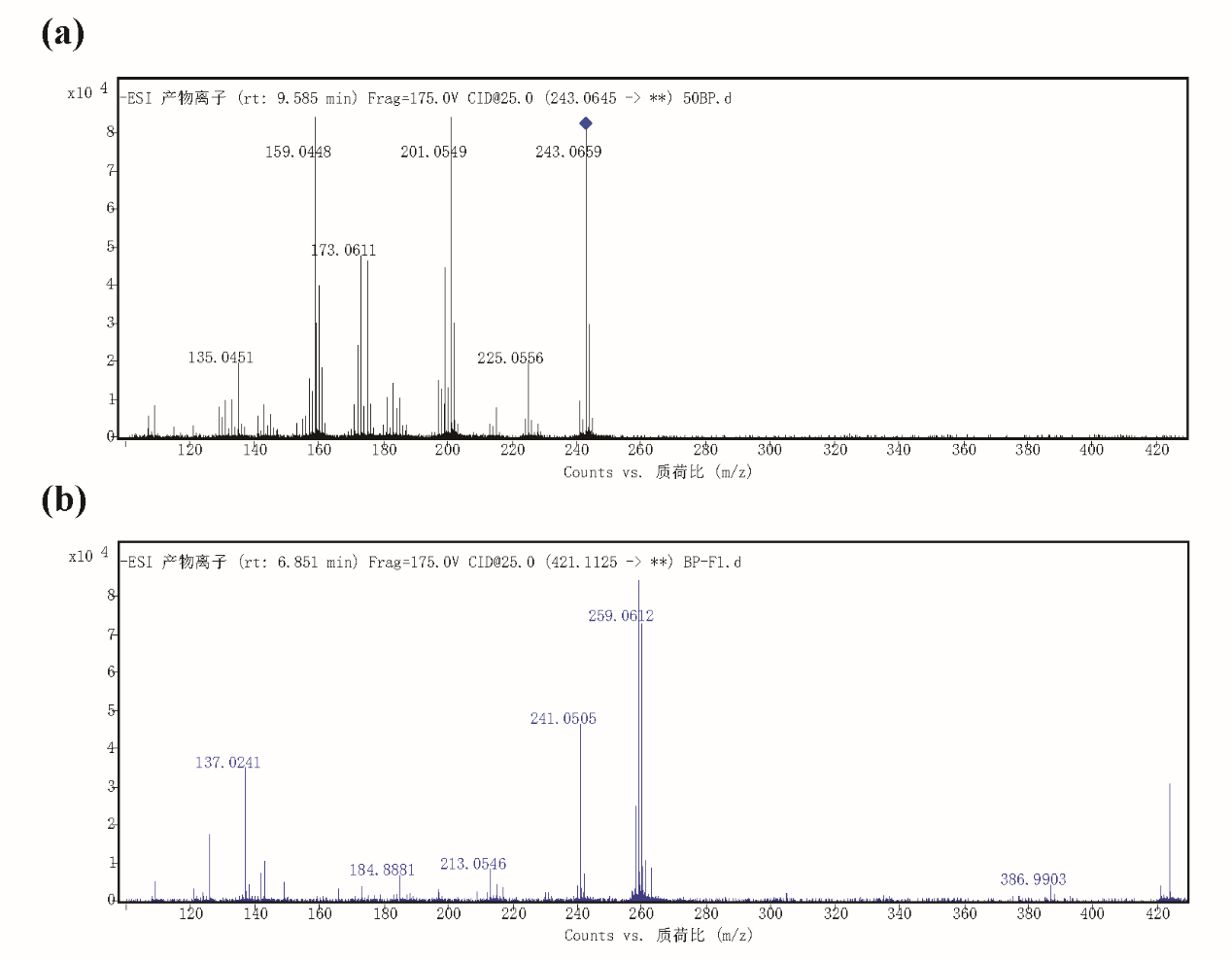

**Figure S21. The ESI mass/mass spectra of piceatannol (a) and its hydroxylated glycosylated products (b).**

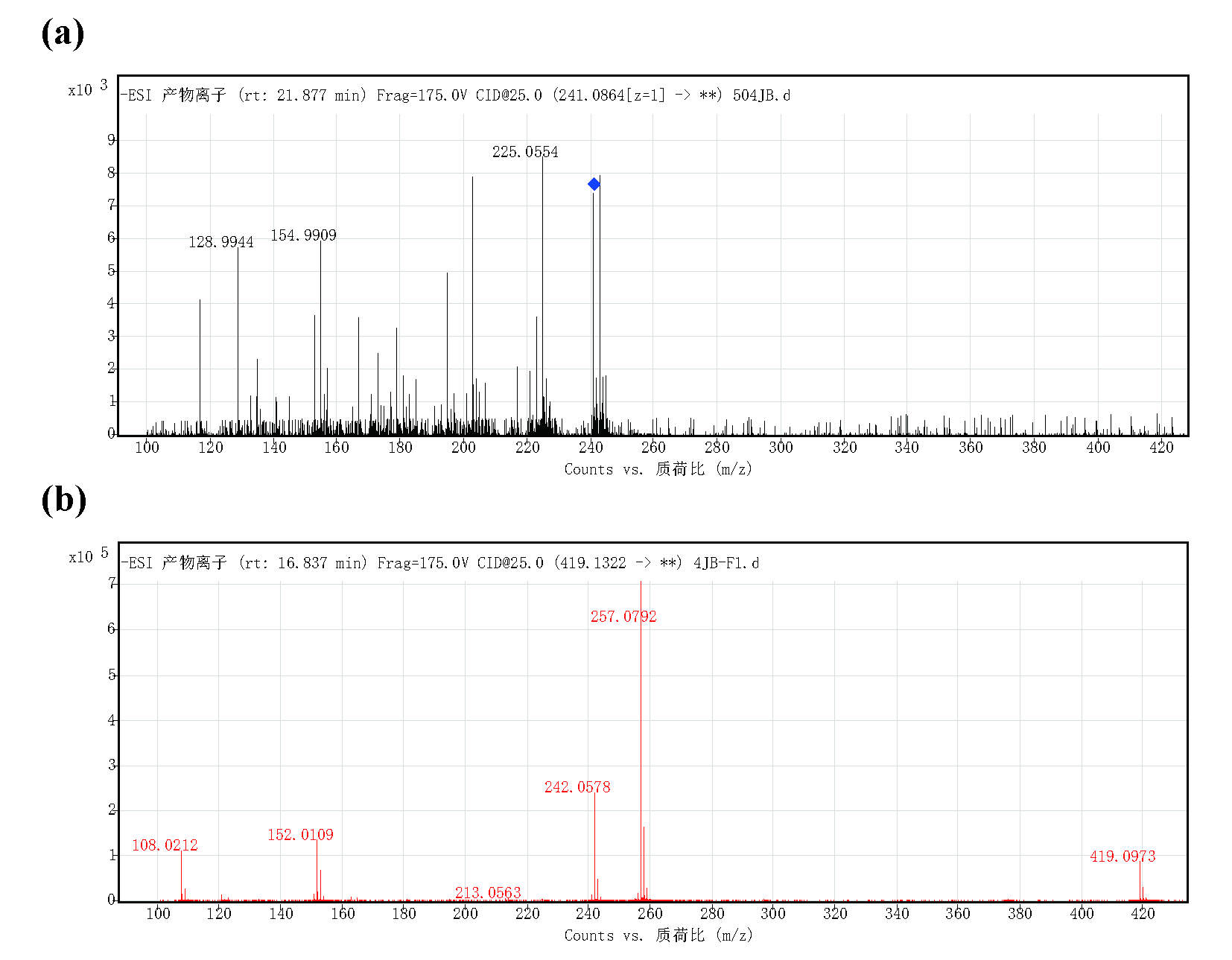

**Figure S22. The ESI mass/mass spectra of desoxyrhapontigenin (a) and its hydroxylated glycosylated products (b).**

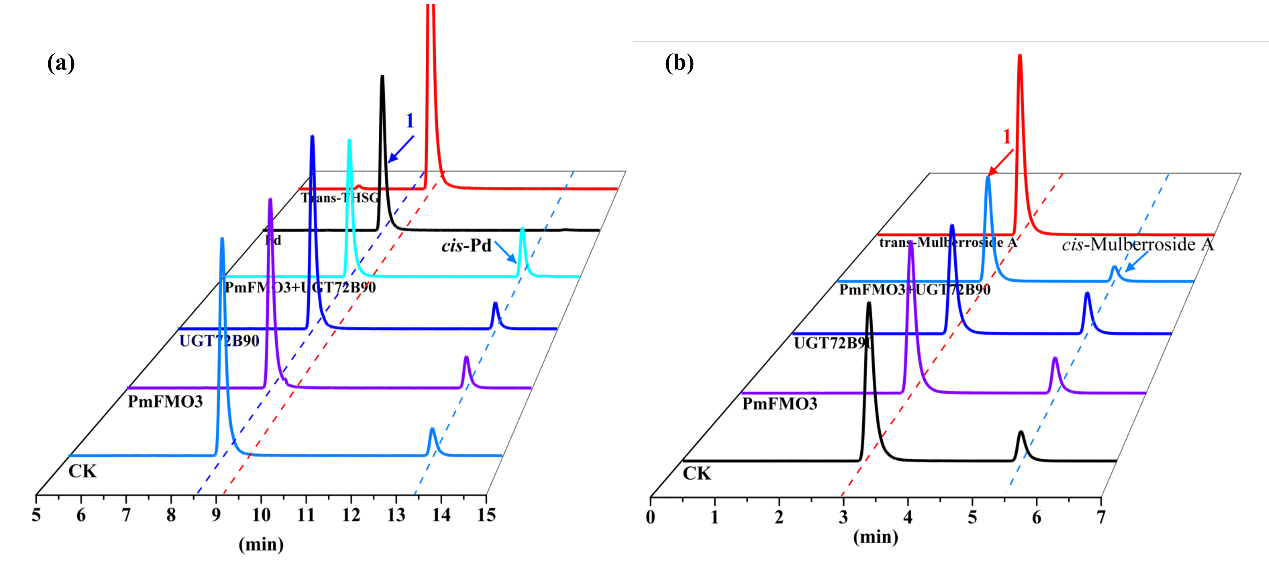
**Fig. S23. The confirmation of the substrate selectivity of PmFMO3 for piceid and mulberroside A by *in vitro* enzymatic reaction. (a)** The multiple reaction monitoring (MRM) chromatograms which detecting the *in vitro* enzymatic reaction product when piceid was added. THSG cannot be detected in any *in vitro* reaction. **(b)** The multiple reaction monitoring (MRM) chromatograms which detecting the *in vitro* enzymatic reaction product when mulberroside A was added. the catalysate same as of 4 cannot be detected in any *in vitro* reaction.

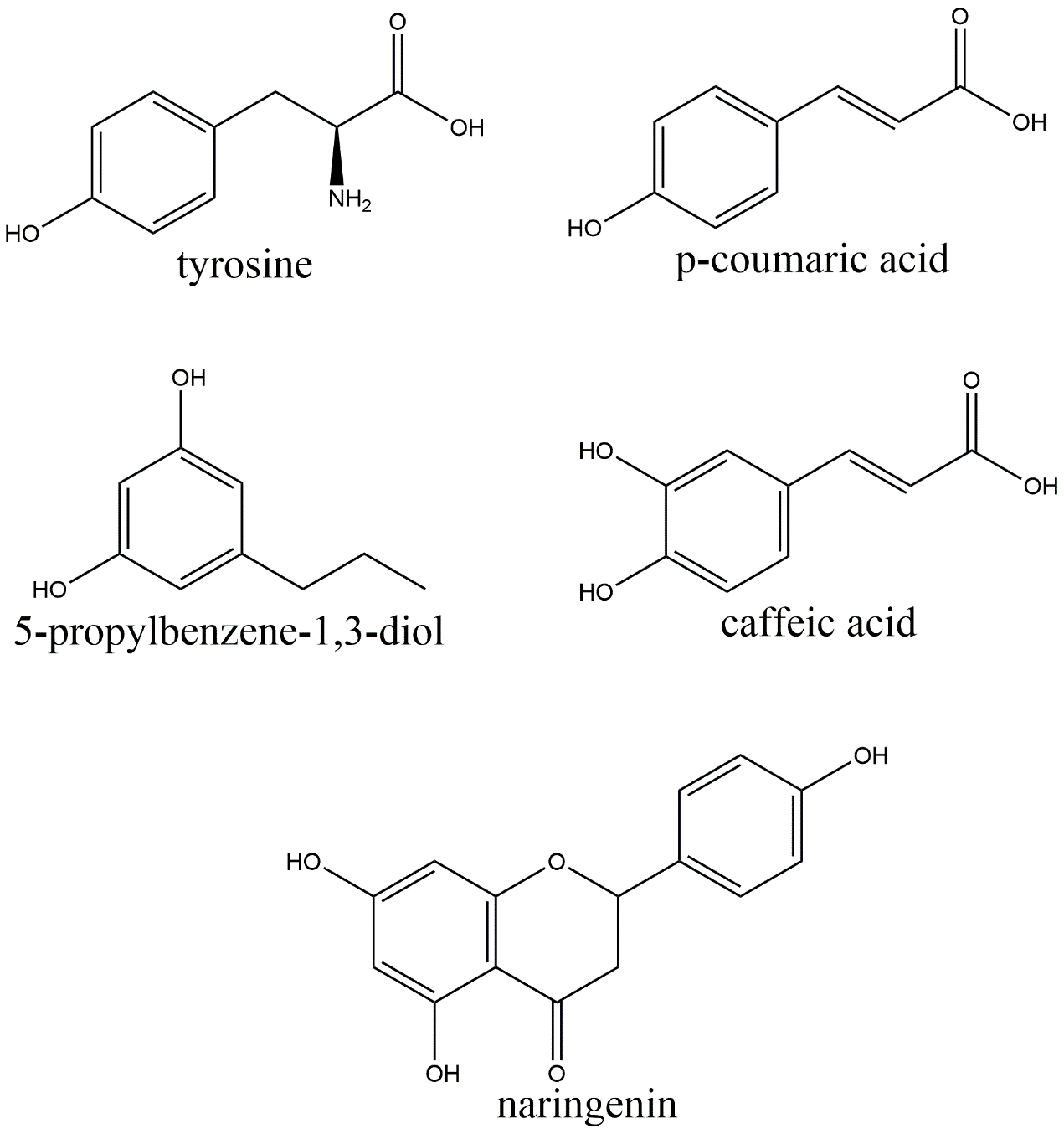

**Fig. S24. The chemical structural formula of other type compounds with the similar structure used for detecting the substrate selectivity of PmFMO3.**

**Supplementary Methods**

**Genome size estimation**

Genome size was estimated by flow cytometry and K-mer frequency distribution analysis. For flow cytometry, *Solanum lycopersicum*, which the genome size has been identified as approximately 900M by genome sequencing was used as internal reference standard (Sato et al., 2012; Li et al., 2023). About 20 mg of *P. multiflorus* and *S. lycopersicum* young leaves were chopped and soaked in 1 mL LB01 buffer (LOUREIRO et al., 2006). The cell culture was collected and filtered through a 400 mesh nylon strainer. The samples were stained with 100 μg mL^−1^ PI simultaneously with 100 μg mL^−1^ RNase in an ice bath for 10 min. The nuclear fluorescence was measured by the MoFlo XDP high-speed flow cytometer (Beckman Coulter Inc., USA) with a 70 μm ceramic nozzle at 60 psi sheath pressure. PI fluorescence with a solid-state laser (488 nm) and detected with a 625 nm HQ band-pass filter. The FL3-Height\SSC-Height gate method was used to eliminate debris, cell fragments, and dead cells. Single cell and double cells were discriminated by using FL3-Height/FL3-Area. For K-mer frequency distribution analysis, a 350 bp paired-end library was constructed and sequenced on the Illumina Novaseq 6000 platform by Berry Genomics Company. After filtered the adaptors and low-quality reads, the clean reads were used to estimate the genome size and heterozygosity. K-mer distribution was estimated by jellyfish (Marçais and Kingsford, 2011). The genome size and heterozygosity ratio were estimated by the online tool of GenomeScope (Ranallo-Benavidez et al., 2020).

**Genome sequencing**

For circular consensus sequencing of *P. multiflorus*, high-quality genomic DNA was isolated from young leaves of *P. multiflorus* by the CTAB method. Two 15 kb DNA SMRTbell library were constructed and sequenced using a single 8 M SMAT Cell on the PacBio Sequel II platform (Pacific Biosciences, CA, USA). The CCS reads were generated by SMRTLink 9.0 software.

To construct the Hi-C library, young leaves of *P. multiflorus* were fixed in 1% formaldehyde for crosslinking. DNA was digested by the *Dpn* II restriction enzyme. The DNA ends were filled and labeled with biotin. The resulting blunt-end fragments were then ligated, purified and random sheared into 300-500 bp fragments. After quality control test, 150 bp PE sequencing of the Hi-C library were performed on the Illumina Novaseq 6000 platform by Berry Genomics Company.

**Genome assembly**

To primarily assemble the genome of *P. multiflorus*, a total of 742.34 Gb of raw PacBio subreads were filtered and corrected using the PacBio circular consensus sequencing (pbccs) pipeline with default parameters. The resulting CCS reads were subjected to hifiasm (Cheng et al., 2021) for *de novo* assembly with default parameters. We corrected the primary contigs with the Pilon program (Walker et al., 2014) with default parameters using Illumina paired-end reads. Then, the redundancy of the assembled sequences was removed to improve the continuity of the assembled contigs using Redundans (Pryszcz and Gabaldón, 2016) and Purge Haplotigs (Roach et al., 2018).

**Chromosomal genome construction**

The paired-end Hi-C reads were uniquely mapped onto the contigs using Juicer(Durand et al., 2016b), and the non-duplicate mapped results were used as the input for the 3D-DNA pipeline (Dudchenko et al., 2017) to construct the genome sequence. Juicebox(Durand et al., 2016a) was used to fine-tune the assembled genome in a graphic and interaction matrix. To assess the completeness of *P. multiflorus* genome, the error rate considering the homozygous mutation was estimated by mapping the 106.44 Gb whole-genome sequence (WGS) reads onto this assembly by BWA software (Li and Durbin, 2010).

**Repeat sequence annotation**

Repeat structures were analyzed by a combined strategy of *de novo* prediction and homology-based prediction. A *de novo* repeat library of the *P. multiflorus* genome was built by RepeatModeler. Using this library, we processed repetitive sequences to annotate, classify, and mark by RepeatMasker (Tarailo-Graovac and Chen, 2009). Two built libraries were combined with the Repbase (Bao et al., 2015) and Dfam (Hubley et al., 2016) databases with default parameters. Simple sequence repeats (SSRs) were identified using MISA (Thiel et al., 2003). Long terminal repeats (LTRs) were identified by LTR_retriever (Ou and Jiang, 2018) according to the suggested method. The insertion time was estimated using the formula T = Ks/2r where Ks is the divergence rate and r (3.48 × 10^−9^) is the substitution rate.

**Gene model prediction**

Three complementary methods were employed to predict the high-quality protein coding genes. Firstly, Illumina RNA-seq data from three representative tissues (ERT, TcBLC, and ML), were assembled with two different strategies (*de novo* or genome-guided assembly) by Trinity(Haas et al., 2013). The assembled RNA-seq data were then aligned to the assembled genome and its evidence-based prediction was performed using PASA(Haas et al., 2003). Secondly, the ab initio methods AUGUSTUS(Stanke et al., 2006), SNAP(Korf, 2004), and GeneMarkHMM (Lukashin and Borodovsky, 1998) with default parameters were used to predict gene models with training of the best candidate genes obtained from PASA (Haas et al., 2003). Thirdly, protein sequences from closely related species downloaded from NCBI and other bioinformatics datebases, were used to annotate protein homologs of *P. multiflora* by GenomeThreader. Finally, the annotation results generated from evidence-based prediction, ab initio prediction, and homologous mapping were combined by EVM (Haas et al., 2008) to integrate the consensus gene model, and genes were renamed according to their position in the genome sequence with the prefix Pm (*P. multiflorus* genome). Compared with the eudicotyledons_odb10 database, the single-copy orthologs in the assembled genome were identified and the completeness of the assembly was evaluated using Benchmarking Universal Single-Copy Orthologs (BUSCO) (Simão et al., 2015).

**Noncoding and protein-coding RNAs functional** **annotation**

Noncoding RNAs and small RNAs were annotated by alignment to the Rfam and miRNA databases using INFERNAL (Nawrocki et al., 2009) and BLASTN with default parameters, respectively. The tRNA genes were identified using the tRNAscan-SE software (Lowe and Eddy, 1997) with default parameters. rRNA fragments were predicted by alignment to rRNA sequences based on BLASTN analysis (*E*-value of 1e-10). Functional annotations of the predicted protein-coding genes were performed (*E*-value of 1e−5) against publicly available protein databases including eggNOG, GO, KEGG, NR, SwissProt databases. Then, GO terms were assigned by the Blast2GO pipeline (v 3.1.3) (Conesa et al., 2005) with default parameters. GO enrichment and KEGG pathway analysis were performed by Gene Ontology (Ashburner et al., 2000) and KEGG datebase (Kanehisa, 2002).
